# Transgenerational pathogen defence extends beyond behavioural avoidance

**DOI:** 10.64898/2026.04.10.717426

**Authors:** Martyna K. Zwoinska, Anasthasia Nathania Widjaja, Aksel Deniz Akgül, Ayşe Sara Altan, Doğukan Aydın, Duygu Çukurbağlı, Elizabeth M. L. Duxbury, Isaac Harris, Alexei A. Maklakov, Nikos Tsardakas Renhuldt, Adithya Ganesh Venkataramani, Martin I. Lind, Hwei-yen Chen

**Author notes:** Corresponding authors: Martyna K. Zwoinska and Hwei-yen Chen.

## Abstract

Transgenerational plasticity can prepare descendants for environmental stress, but its generality and adaptive value remain unclear. Here we tested for adaptive transgenerational effects of exposure to the bacterial pathogen *Pseudomonas vranovensis* across five *Caenorhabditis* species. Pathogen exposure reduced rate-sensitive fitness in all five species, but behavioural responses varied across species and generations. In *C. elegans*, exposed parents and their F2 descendants showed increased pathogen avoidance. In *C. remanei*, pathogen exposure increased attraction in females, whereas their F2 female descendants showed greater avoidance. We found no F2 behavioural response in *C. kamaaina*, *C. tropicalis* or *C. briggsae*. Remarkably, however, ancestral exposure increased F2 larval survival on the pathogen in *C. elegans*, *C. remanei* and *C. tropicalis*. In *C. tropicalis* this survival benefit occurred without behavioural avoidance. The two species showing transgenerational avoidance had the closest sequence matches between the bacterial small RNA Pv1 and its candidate neuronal target *maco-1*, suggesting a possible molecular basis for variation in the response. Our results show that shared susceptibility to a pathogen does not imply similarity in transgenerational response, and that ancestral exposure can benefit descendants without inheritance of pathogen avoidance.

## Introduction

Transgenerational epigenetic inheritance (TEI) is a form of phenotypic plasticity in which environmentally induced effects persist in descendants that have not experienced the environmental conditions that triggered the effect. TEI can be mediated by several epigenetic mechanisms, including small RNAs, chromatin modifications and DNA methylation.

Transgenerational effects have been reported in response to a wide range of biotic and abiotic factors, including pathogens, predators, temperature and starvation^1,2^. The resulting phenotypic changes are diverse and can involve different responses to environmental stress^3^. TEI may provide a rapid and reversible response to environmental change, operating on timescales between within-generation plasticity and genetic adaptation^4,5^. Theory predicts that it can be favoured when parental environments reliably predict the conditions descendants will encounter^5,6^. However, when parental cues are unreliable or inherited responses carry greater costs than benefits, transgenerational responses may be neutral or maladaptive^5,7^. However, the generality of transgenerational effects remains elusive, with effects varying across taxa, traits and environmental contexts^1,2^. Comparative studies across closely related species are therefore needed to shed light on whether and when transgenerational responses are adaptive.

An especially striking example of TEI occurs in the nematode worm *Caenorhabditis elegans*, which can learn to avoid pathogenic bacteria and transmit this learned avoidance to naïve descendants for up to four generations^8–10^. Studies of this response have focused on *Pseudomonas* bacteria, which are common members of the worm’s bacterial community^11^.

While many *Pseudomonas* bacteria serve as nutritious food sources^11^, *P. aeruginosa* strain PA14^10,12^, *P. vranovensis* strain GRb0427^9^ and *P. fluorescens* strain PF15^8^ are pathogenic to *C. elegans*. Despite this danger, naïve worms are often attracted to these pathogens and learn to avoid them only after ingestion. Following parental learning, descendants that have never encountered the pathogen can inherit this avoidance, reducing their otherwise elevated risk of exposure and infection^8–10^.

The molecular basis of this response has been characterised in considerable detail. Both learning and TEI of pathogen avoidance in *C. elegans* are mediated by bacterial small RNAs processed through the worm’s RNA interference (RNAi) machinery, a gene-silencing pathway that acts on double-stranded RNA (dsRNA)^8,9,13^. These bacterial small RNAs include P11 (*P. aeruginosa*), Pv1 (*P. vranovensis*) and Pfs1 (*P. fluorescens*). They exhibit 16-to 17-nucleotide homology to two neuronal genes in *C. elegans*: *maco-1*, a neuronal homologue of the human nervous-system-specific ER membrane protein Macoilin and the target of P11 and Pv1, and *vab-1*, which encodes an ephrin receptor upstream of *maco-1* and is the target of Pfs1. Upon ingestion, bacterial small RNAs are taken up via the dsRNA transporter SID-2 and processed in the intestine by the DCR-1 (Dicer) endoribonuclease. They are then transported to the germline through the systemic RNAi channel SID-1 and subsequently from the germline to neurons via *cer-1* retrotransposon-encoded virus-like particles^14^. This leads to down-regulation of *vab-1* and *maco-1*, which in turn leads to up-regulation of *daf-7* in the ASI neurons and thereby the behavioural change from attraction to avoidance^8,9,13^.

The extent to which pathogen-induced TEI and its underlying molecular mechanisms extend beyond *C. elegans* remains unclear. *Caenorhabditis* species are genetically diverse and occupy distinct ecological niches, exposing them to different microbial communities and potentially favouring different defence strategies^15–17^. Even when confronted with the same pathogen, species may therefore differ in their susceptibility and in the responses induced by parental exposure. Consistent with this, *C. elegans*, *C. kamaaina*, *C. briggsae* and *C. tropicalis* differ in both their susceptibility and intergenerational (F1) responses to *P. vranovensis* strain BIGb0446^18^. However, whether these intergenerational effects persist transgenerationally (F2+) across *Caenorhabditis* remains largely unexplored. Even within *C. elegans*, studies have reported inconsistent inheritance of learned pathogen avoidance^19–21^, raising questions about the reproducibility of the response. The ecological relevance of the pathogen challenge is also important when evaluating adaptive significance. *P. vranovensis* GRb0427 was isolated from the natural microbiota of *C. elegans*^11^, whereas several commonly used pathogens, including the clinical isolate *P. aeruginosa* PA14, *Salmonella enterica* Typhimurium MST1 and *P. aeruginosa* PAO1, have less clear ecological relevance^9,22,23^. Together, these gaps leave unresolved how general pathogen-induced transgenerational effect is across *Caenorhabditis* and whether its consequences are adaptive.

Here, we used *P. vranovensis* GRb0427 to compare pathogen-induced transgenerational responses across five *Caenorhabditis* species representing independent branches of the Elegans group^24^: *C. kamaaina* QG122, *C. elegans* N2, *C. tropicalis* JU1373, *C. remanei* PB4641 and *C. briggsae* AF16. We first tested whether *P. vranovensis* reduced fitness across species and examined whether candidate components of the *C. elegans* TEI pathway were conserved. We then established the reproducibility of learned avoidance in *C. elegans* and used a standard protocol to compare within-and transgenerational behavioural responses across species and sexes. Finally, we tested whether parental exposure to *P. vranovensis* affected descendant survival upon subsequent pathogen challenge. Together, these experiments allowed us to assess both the generality and potential adaptive value of pathogen-induced transgenerational responses across *Caenorhabditis*.

## Results

### P. vranovensis is pathogenic to Caenorhabditis worms

We first tested whether *P. vranovensis* strain GRb0427 is pathogenic across the five *Caenorhabditis* species included. Age-synchronised worms were raised to the pre-adult stage (L4) and then transferred to plates seeded with either *E. coli* or *P. vranovensis* (hereafter E-trained worms and P-trained worms, respectively; Figure 1). After 24 hours of training on either bacterial treatment, adult hermaphrodites or inseminated females were washed and transferred to

**Figure 1:**
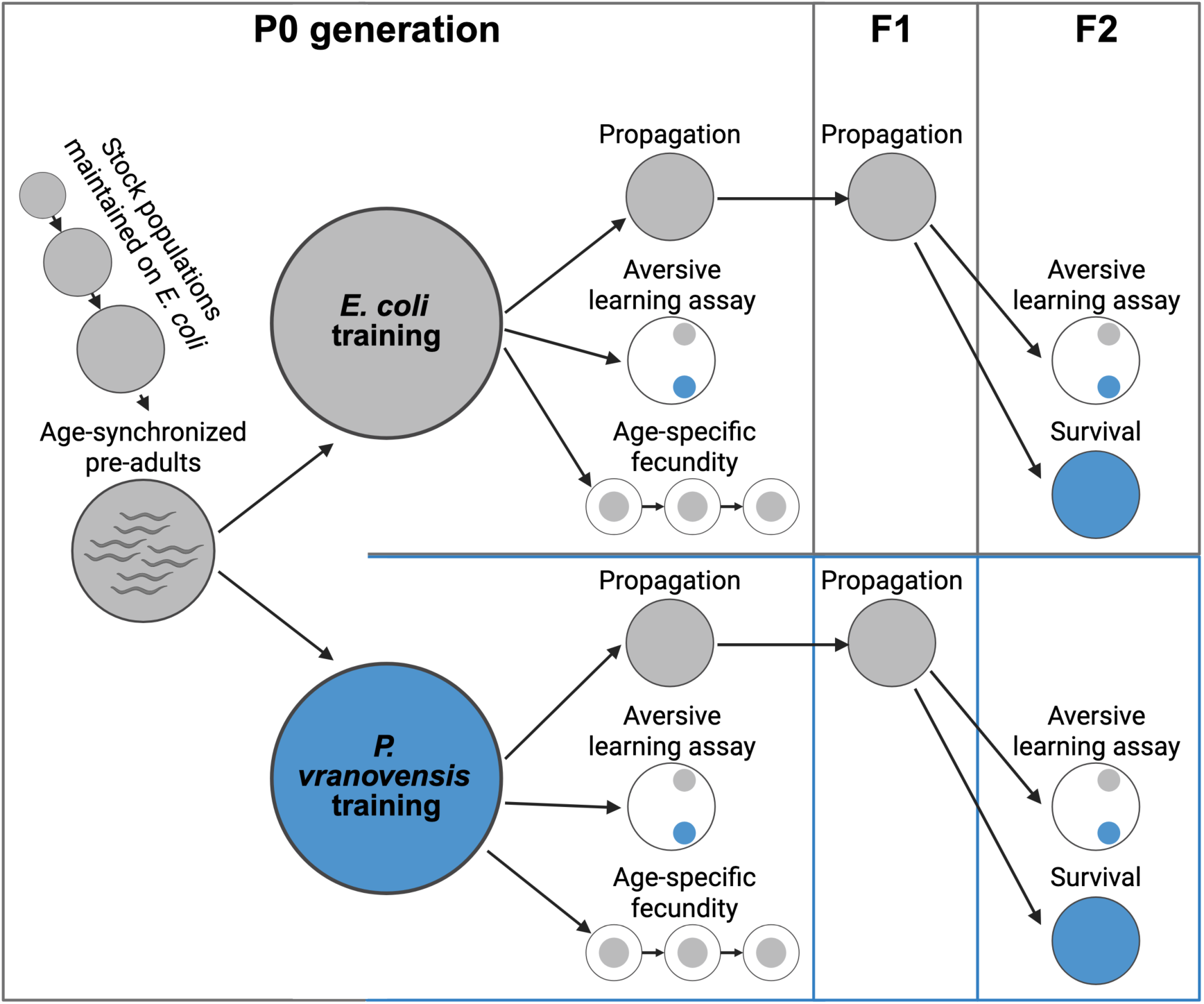
Procedures for bacterial training on *E. coli* (benign food; grey) or *P. vranovensis* (pathogen; blue), aversive learning, transgenerational behavioural assays, age-specific fecundity and survival assays. In the P0 parental generation, age-synchronised pre-adults were randomly assigned to either E-or P-training regimes. Following bacterial training, P0 worms were assessed for aversive learning and age-specific fecundity. F1 worms were propagated on *E. coli*. For the F2 transgenerational behavioural assay, F2 worms were propagated on *E. coli*; for the survival assay, F2 embryos were placed directly on *P. vranovensis*. In the F2 generation, worms underwent choice assays and pathogen survival assays.

*E. coli*-seeded NGM plates

Worms were left to lay eggs and transferred to fresh plates every 24 hours. Age-specific fecundity was measured as the number of offspring larvae produced each day.

For each species, we analysed the effect of bacterial training on age-specific fecundity as well as an individual-based, rate-sensitive estimate of fitness (individual fitness, **λ**_ind_). In particular, **λ**_ind_ combines age-specific fertility, survival and reproductive timing over the assay period^25^: for individual *i*, **λ**_ind_ is the largest root of **Σ***f_x_l_x_***λ**_ind_^−x^ = 1, where *f_x_* is fertility at age *x* and *l_x_* is survival at age *x* (either 1 or 0). We calculated **λ**_ind_ using the *lambda* function in the R package popbio.

Since *Caenorhabditis* nematodes have boom-and-bust population dynamics in nature^26^, early fecundity is crucial for rapid expansion on ephemeral food sources.

We confirmed that *P. vranovensis* strain GRb0427 reduced **λ**_ind_ in all five species tested: *C. kamaaina*, *C. elegans*, *C. tropicalis*, *C. remanei* and *C. briggsae*, with large effect sizes (two-sample t-tests: Table 1). *P. vranovensis* training also significantly reduced age-specific fecundity in *C. kamaaina* and *C. briggsae* (generalised linear mixed-effects models, GLMMs: *C. kamaaina*: p = 0.006; *C. briggsae*: p < 0.001; Figure 2; Table S1), but not in *C. elegans*, *C. tropicalis*, and *C. remanei* (*C. elegans*: p = 0.362; *C. tropicalis*: p = 0.808; *C. remanei*: p = 0.126; Figure 2; Table S1).

**Figure 2:**
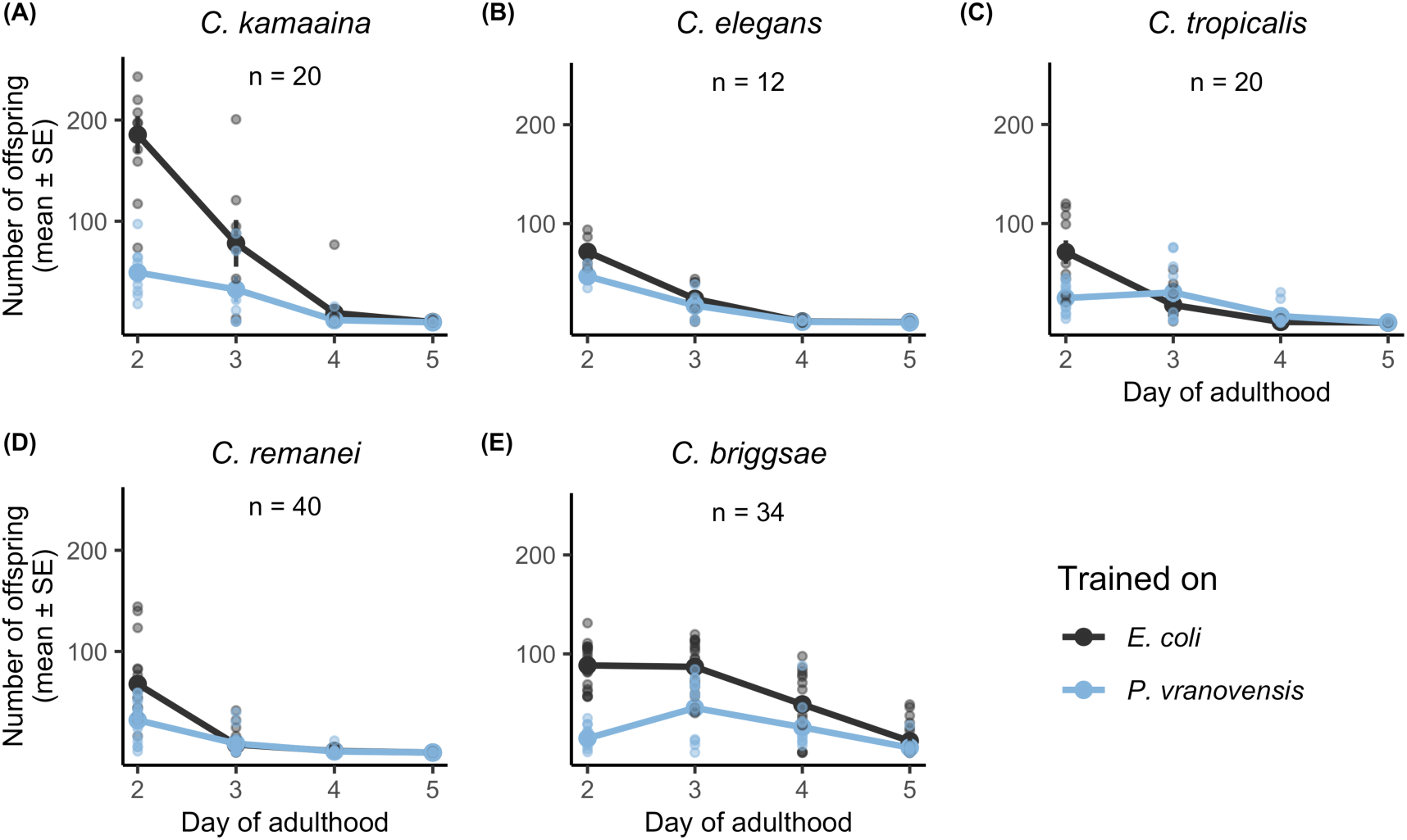
Age-specific fecundity of *C. kamaaina* (A), *C. elegans* (B), *C. tropicalis* (C), *C. remanei* (D) and *C. briggsae* (E) females or hermaphrodites following 24 h of training on *E. coli* (grey symbols) or *P. vranovensis* (blue symbols). Fecundity was measured on *E. coli*.

**Table 1:**
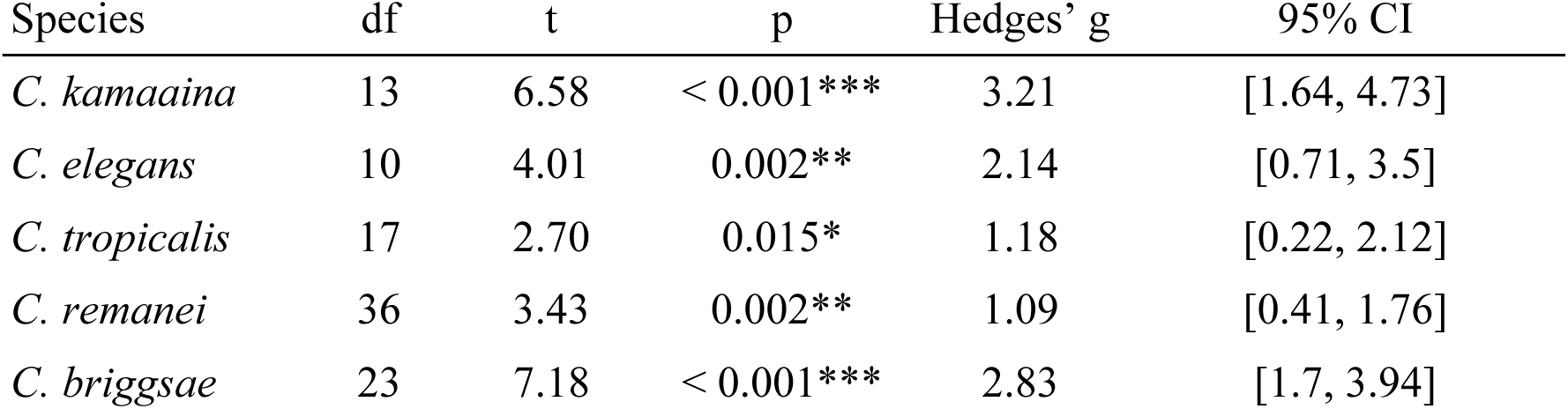
t-test statistics, effect sizes (Hedges’ g) and 95% confidence intervals (CI) for the effect of bacterial training (E-versus P-training) on rate-sensitive individual fitness (**λ**_ind_) in the P0 generation across the five species. df: degrees of freedom.

Although bacterial training had no significant main effect on age-specific fecundity in *C. elegans*, *C. tropicalis* or *C. remanei*, P-trained worms produced fewer offspring on the first assay day (the second day of adulthood). In *C. tropicalis* and *C. briggsae*, the effect of training changed significantly with age (training × age interaction: *C. tropicalis*: p = 0.007; *C. briggsae*: p = 0.025; Table S1), while no significant training × age interaction was detected in *C. elegans* or *C. remanei*. Because **λ**_ind_ gives greater weight to early reproduction, the reduction in early fecundity likely contributed to the lower **λ**_ind_ observed in these species.

Training on *P. vranovensis* significantly reduced age-specific fecundity in *C. kamaaina* and *C. briggsae* (see Table S1 for more details). Solid points: mean fecundity; error bars: standard errors; translucent points: raw data. n indicates the number of plates included for each species.

### Components of the *C. elegans* TEI pathway vary across *Caenorhabditis*

We investigated candidate components of the *C. elegans* TEI pathway, specifically Pv1-*maco-1* sequence homology and the presence of a putatively active *cer-1* retrotransposon. A previous study showed that a four-nucleotide mismatch is sufficient to abolish the inheritance of learned avoidance^9^, so we considered sequences with up to three mismatches. Of the five species, only *C. elegans maco-1* showed perfect 16-nucleotide homology with the bacterial small RNA Pv1 (Figure 3). *C. remanei* showed a single-nucleotide mismatch, whereas *C. briggsae* showed three mismatches (Figure 3). No *maco-1* sequence with ≤3 mismatches to Pv1 was identified in *C. kamaaina* or *C. tropicalis* (Figure 3).

**Figure 3:**
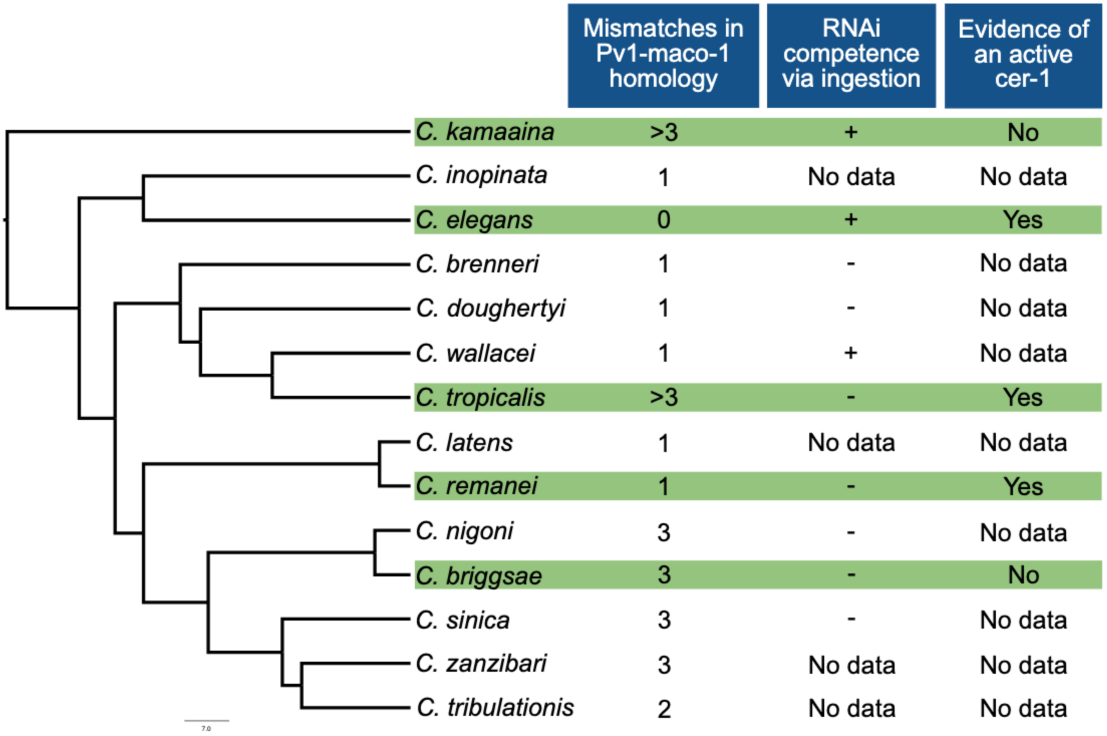
Phylogeny, number of mismatches in Pv1-*maco*-1 sequence homology, evidence of a putatively active *cer*-1 retrotransposon and sensitivity to ingested dsRNA across species in the *Elegans* group. Species highlighted in green were tested in this study. The number of mismatches represents nucleotide differences in the 16-nt Pv1-*maco*-1 homologous sequence (see Table S6 for details). Sensitivity to ingested dsRNA: data from Nuez and Felix^27^; +, sensitive; -, insensitive; no data: not tested. Phylogeny adapted from Fusca et al.^24^.

We found evidence of a putatively active *cer-1* retrotransposon in *C. tropicalis and C. remanei* (Figure 3; Figure S1; Supplemental files S1-S3). In *C. tropicalis,* we detected 17 polymerase (POL) hits, including seven near full-length POL regions (>= 90% of the *C. elegans* Cer-1 POL) on chromosomes I, II and X. Ten of these POL regions lay within elements bounded by two terminal repeats (LTRs), and six of these elements also contain the *cer-1* primer-binding site (PBS). *C. remanei* carried the greatest number of *cer-1* like copies among the four species analysed, with 54 POL hits, including 22 near full-length POL regions on chromosomes II-V, X and four placed on unplaced scaffolds, and 29 LTR elements, 13 of which also contained both a POL region and a PBS. In both species, the LTRs of most elements were <= 99% identical (9 of 10 in *C. tropicalis*, 18 of 29 in *C. remanei*), suggesting recent insertion. GAG regions, which encode proteins needed for virus-like particle (VLP) formation, were also present in most of these elements (9 of 10 and 19 of 29, respectively). By contrast, there was weak evidence for active *cer-1* elements in *C. kamaaina* and *C. briggsae* (Figure 3; Figure S1; Supplemental files S1-S3). *C. kamaaina* had 12 POL hits, only one of which was full length. None lay within LTRs, and no PBS was detected. GAG homology was detected at 2 sites. However, the heavily fragmented assembly (8,860 scaffolds) of *C. kamaaina* limited our detection power. *C. briggsae* only had four POL hits, none of which were full length. A single LTR copy with a PBS was located on chromosome III, but its POL region was incomplete (83% of the *C. elegans* POL), and its LTRs were less similar than *C. tropicalis* or *C. remanei*, consistent with degraded or older copies.

A previous study showed that sensitivity to ingested double-stranded RNA varies across *Caenorhabditis* worms^27^. Of the five species included in our study, *C. elegans* and *C. kamaaina* are sensitive to ingested RNA, but not *C. tropicalis*, *C. remanei*, and *C. briggsae*^27^ (Figure 3). Together, these results show that candidate components of the *C. elegans* Pv1-mediated TEI pathway are not conserved across the *Elegans* group.

### Reproducibility of learned avoidance in *C. elegans*

Much of the debate surrounding the reproducibility of TEI of pathogen avoidance comes from the inclusion of the anaesthetic sodium azide during assays that measure worms’ attraction or avoidance to the pathogens^19–21^. In a standard chemotaxis assay, a test compound is placed at one edge of an agar plate and a control solvent at the opposite edge. Worms are released from an origin point equidistant from both locations and allowed a fixed period of time to migrate.

Sodium azide is commonly used to immobilise worms once they reach either side, thereby preserving their first choice. Of particular concern is that naïve attraction to *Pseudomonas* pathogens may disappear if worms leave the pathogen after experiencing harm. Nonetheless, the assay period is typically short (one hour), whereas behavioural changes generally require longer exposure (>4 hours)^10,28^. More importantly, worms migrate freely in their natural habitats. If worms can avoid harm by instantaneously leaving the pathogens, it raises the question of why they would evolve a multigenerational response.

We tested the reproducibility of the experimental paradigm in *C. elegans*. Specifically, we set up independent experimental blocks of aversive learning assays in which worms were either allowed to migrate freely (i.e., no anaesthetics) or were immobilised upon arriving at their first choice. We closely followed a previously published protocol^29^ with minor modifications (Figure 1; see also Method details). The most notable change was the use of potassium azide instead of sodium azide due to local chemical safety regulations. However, both chemicals have a similar immobilising effect^30^. We focused on within-generation behavioural responses to *P. vranovensis*. All behavioural assays were analysed using the standard ANOVA approach to allow for comparison with previous studies^9^; we report these results in the main text (see below). In addition, we complemented our analyses with Wilcoxon tests, t-tests, and generalised linear models (GLMs) to ensure robustness to deviations from model assumptions (see Table S2); these results are reported in Supplemental Figures and Tables.

We tested worms’ food choice after 24 hours of training on either *E. coli* or *P. vranovensis*. E-trained worms showed naïve attraction to *P. vranovensis,* while P-trained worms learned to avoid *P. vranovensis* and preferred *E. coli* (Figure 4; Table 2; Table S2). These results are consistent with previous publications^9^. The direction of the training effect was similar with and without potassium azide (Figure 4; Table 2; Table S2). This suggests that worms do not easily leave their first-choice spot within the 1-hour assay period, and that initial choice accurately reflects their learned or naïve preference.

**Figure 4:**
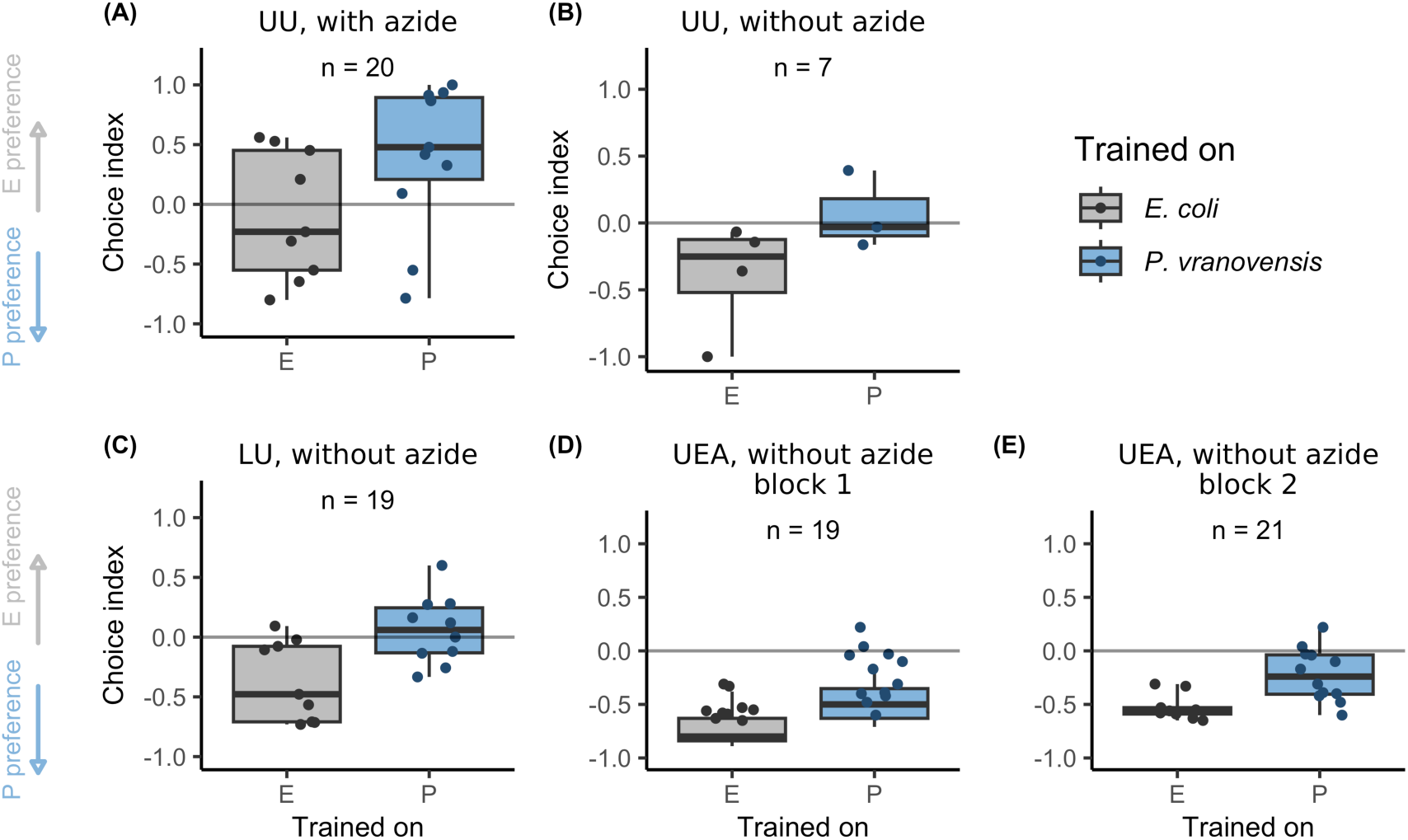
Behavioural responses of *C. elegans* in the P0 parental generation, assessed independently at three laboratories: Uppsala University (UU) and Lund University (LU), Sweden, and the University of East Anglia (UEA), UK. (A) Aversive learning assays performed with potassium azide at Uppsala University. (B) Aversive learning assays performed without potassium azide at Uppsala University. (C) Aversive learning assays performed without potassium azide at Lund University. (D, E) Aversive learning assays performed without potassium azide at the University of East Anglia in two independent experimental blocks (D, block 1; E, block 2). At UU, the effect of training was moderate with azide and large without azide, although neither effect reached statistical significance (see also Table 2 and Table S2). In contrast, the LU and UEA conditions showed large and statistically significant effects (see also Table S2). Note that all data shown in Figures 5 and 6 were collected at Lund University and do not include data shown in (C). Centre line in each box: median; box range: 25th to 75th percentiles; whiskers: minimum-maximum values; dots: raw data from choice-assay plates. Grey boxes and dots represent E-trained worms; blue boxes and dots represent P-trained worms. n indicates the number of plates included in each dataset. Choice index = (# of worms on *E. coli* – # of worms on *P. vranovensis*)/(# of worms on *E. coli* + # of worms on *P. vranovensis*). Positive values indicate preference for *E. coli* (E preference; upward grey arrow) and negative values indicate preference for *P. vranovensis* (P preference; downward blue arrow). The horizontal line at zero indicates no preference

**Table 2:** ANOVA statistics, effect sizes (eta-squared) and 95% confidence intervals (CI) for within-generation response to *P. vranovensis* with or without the inclusion of potassium azide across laboratories (UU, LU and UEA) and experimental blocks.

| Lab | Azide | F-statistic | p | eta-squared | 95% CI |
| --- | --- | --- | --- | --- | --- |
| UU | With azide | $F(1, 18) = 3.72$ | 0.070 | 0.17 | [0, 1] |
| UU | Without azide | $F(1, 5) = 2.55$ | 0.171 | 0.34 | [0, 1] |
| LU | Without azide | $F(1, 17) = 8.98$ | 0.008** | 0.35 | [0.07, 1] |
| UEA | Without azide block 1 | $F(1, 17) = 9.7$ | 0.006** | 0.36 | [0.08, 1] |
| UEA | Without azide block 2 | $F(1, 19) = 11.33$ | 0.003** | 0.37 | [0.1, 1] |

### Within-generation behavioural responses vary across species and sexes

We examined whether *P. vranovensis* training could induce behavioural changes in *Caenorhabditis* species. Since *C. kamaaina* and *C. remanei* are gonochoristic, we analysed each sex separately. *C. elegans*, *C. tropicalis*, and *C. briggsae* are predominantly self-fertilising, with males occurring only rarely^31–33^, and individuals were therefore not sexed in our assays.

Overall, we found substantial variability in within-and transgenerational behavioural responses across species and sexes. Similar variability was also observed across experimental blocks and E-trained baseline responses (see also Table S5, Supplemental Text, and the next section, “Transgenerational behavioural responses vary across species and sexes”). Given this variability, we included complementary statistical approaches in addition to standard ANOVA to assess the robustness of our results. Comparisons between P-and E-trained animals were performed within each generation because experimental handling differed slightly between P0 and F2 (see Supplemental Text).

We found that *C. elegans* showed learned avoidance of *P. vranovensis* after 24 hours of pathogen training (Figure 5), with significant effects and large effect sizes detected by ANOVA (Table 3), a Wilcoxon test, a t-test and a generalised linear mixed-effects model (GLMM; Table S3; see also Supplemental Text and Figure S2), consistent with previous studies^9^. In the other two hermaphroditic species, *C. tropicalis* and *C. briggsae*, no significant behavioural changes were detected (Figure 5; Table 3; Table S3). In the gonochoristic species *C. remanei*, P-trained females showed greater relative preference for *P. vranovensis* than E-trained females after 24 hours of training (Figure 5), with consistent significant effects and moderate-to-large effect sizes across statistical approaches (ANOVA: Table 3; Wilcoxon test, t-test and GLMM: Table S3). By contrast, *C. remanei* males showed no change in preference (Figure 5; Table 3; Table S3; see also Supplemental Text and Figure S4). In *C. kamaaina*, females did not display any detectable behavioural response (Figure 5; Table 3; Table S3; see also Supplemental Text and Figure S6). In contrast, males exhibited a modest behavioural response (avoidance after training): the effect was marginally significant in the ANOVA analysis (Figure 5; Table 3) and significant in complementary statistical approaches with small-to-moderate effect sizes (Figure 5; Table 3; Table S3; see also Supplemental Text and Figure S6).

**Figure 5:**
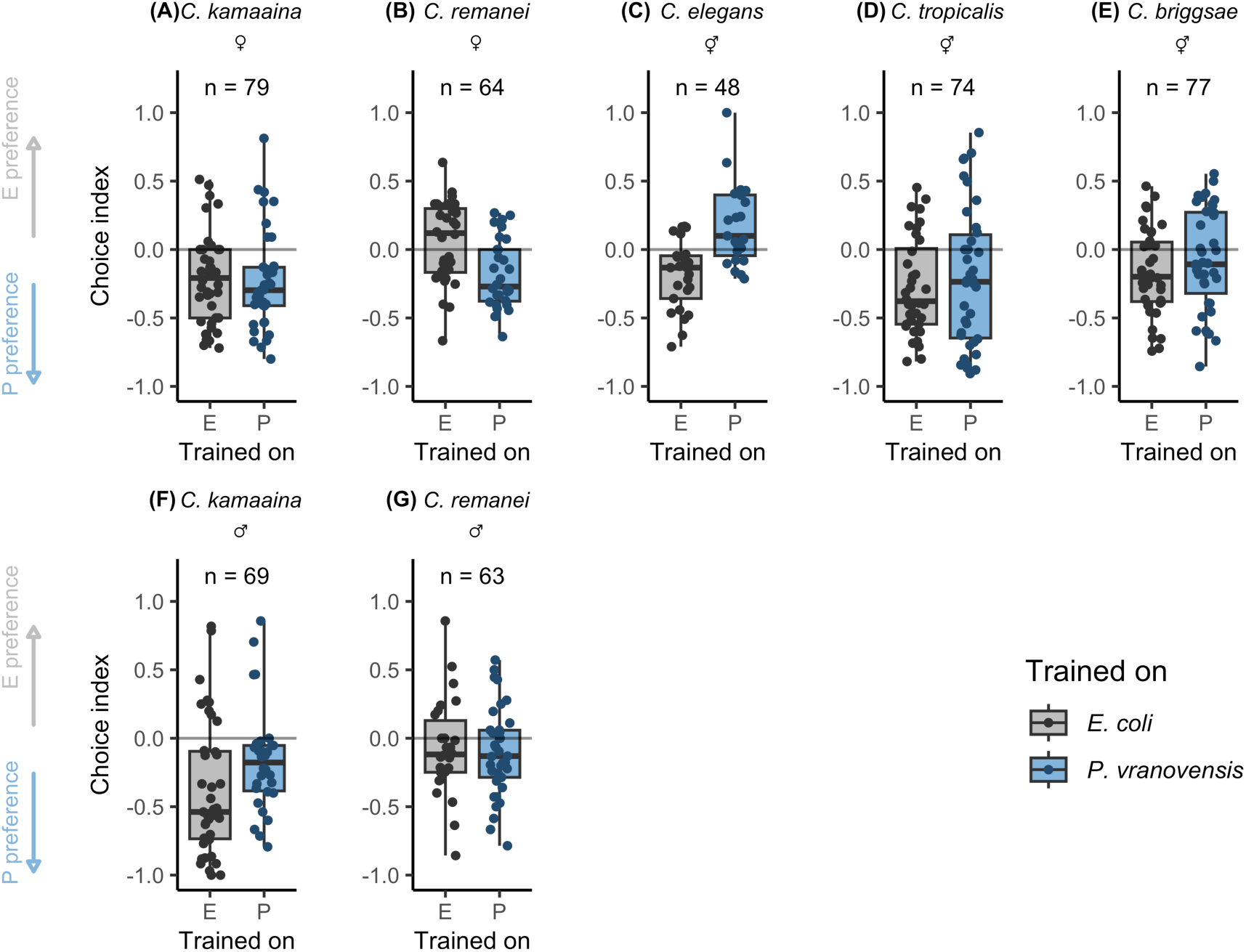
In the P0 parental generation, P-trained *C. elegans* showed greater avoidance of *P. vranovensis* than E-trained controls (C), whereas P-trained *C. remanei* females showed greater relative preference for *P. vranovensis* than E-trained controls (B). Centre line in each box: median; box range: 25th to 75th percentiles; whiskers: minimum-maximum values; dots: raw data from individual choice-assay plates. Grey boxes and dots represent E-trained worms; blue boxes and dots represent P-trained worms. Choice index = (# of worms on *E. coli* – # of worms on *P. vranovensis*)/(# of worms on *E. coli* + # of worms on *P. vranovensis*). n indicates the number of plates included for each species/sex. Positive values indicate preference for *E. coli* (E preference; upward grey arrow) and negative values indicate preference for *P. vranovensis* (P preference; downward blue arrow). The horizontal line at zero indicates no preference.

**Table 3:** ANOVA statistics, effect sizes (eta-squared) and 95% confidence intervals (CI) for within-generation response (P0) to *P. vranovensis* across species and sexes.

| Species | Sex | F-statistic | p | eta-squared | 95% CI |
| --- | --- | --- | --- | --- | --- |
| *C. kamaaina* | Male | F(1, 67) = 3.79 | 0.056 | 0.05 | [0, 1] |
| *C. kamaaina* | Female | F(1, 77) = 0.06 | 0.810 | 0.00 | [0, 1] |
| *C. kamaaina* | Sex-pooled | F(1, 75) = 0.71 | 0.403 | 0.01 | [0, 1] |
| *C. elegans* | Hermaphrodite | F(1, 46) = 22.36 | < 0.001*** | 0.33 | [0.15, 1] |
| *C. tropicalis* | Hermaphrodite | F(1, 72) = 0.45 | 0.505 | 0.01 | [0, 1] |
| *C. remanei* | Male | F(1, 61) = 0.11 | 0.743 | 0.00 | [0, 1] |
| *C. remanei* | Female | F(1, 62) = 12.61 | < 0.001*** | 0.17 | [0.05, 1] |
| *C. remanei* | Sex-pooled | F(1, 61) = 4.3 | 0.042* | 0.07 | [0, 1] |
| *C. briggsae* | Hermaphrodite | F(1, 75) = 1.34 | 0.251 | 0.02 | [0, 1] |

### Transgenerational behavioural responses vary across species and sexes

In the F2 generation, *C. elegans* showed transgenerational pathogen avoidance, consistent with previous findings, with significant effects and large effect sizes across statistical approaches (Figure 6; Table 4; Table S4; see also Supplemental Text and Figure S3). *C. tropicalis* and *C. briggsae* showed no significant F2 behavioural responses (Figure 6; Table 4; Table S4), despite a significant increase in the E-trained baseline from P0 to F2 in *C. tropicalis* (Table S5).

**Figure 6:**
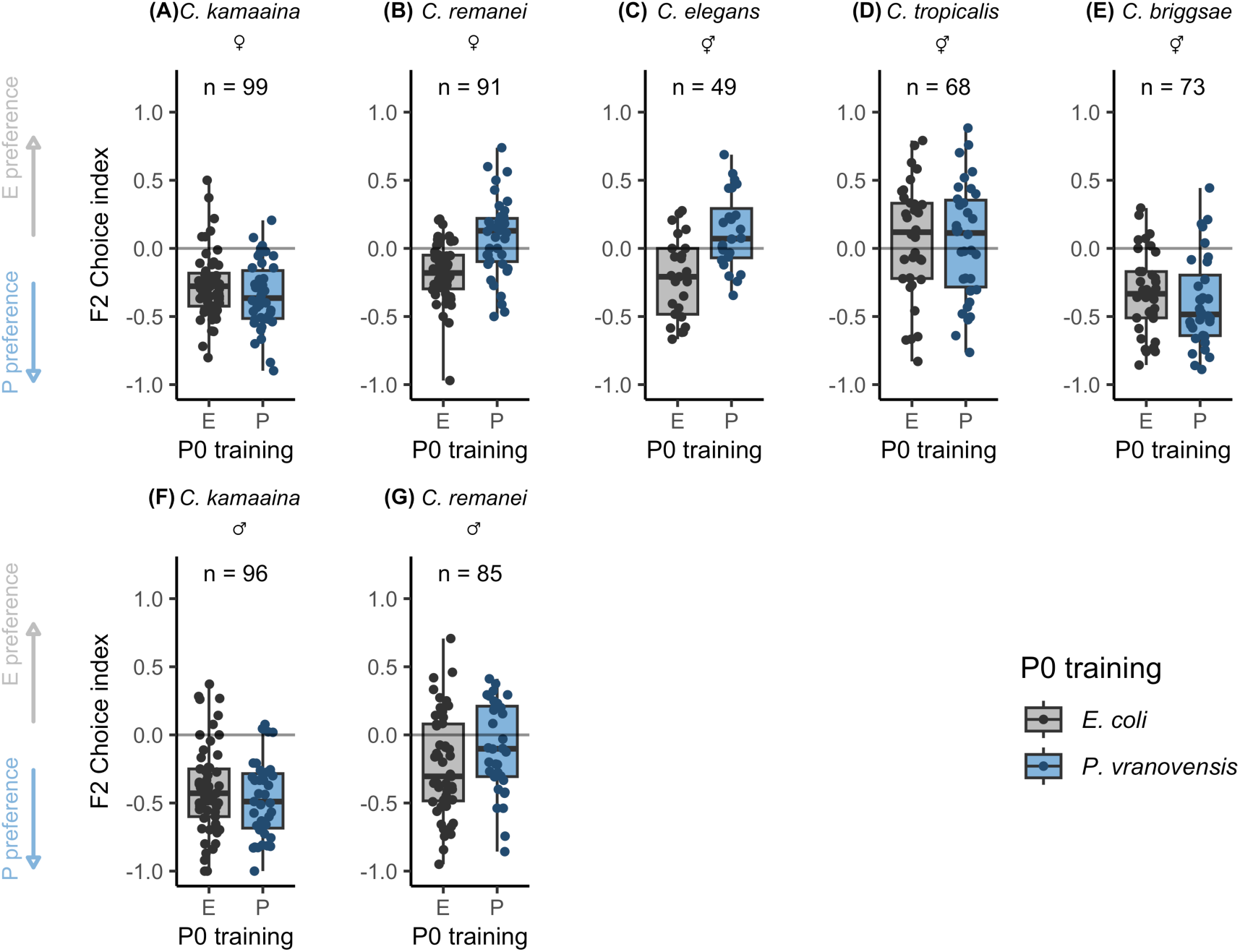
Transgenerational behavioural responses in the F2 generation following parental training on *P. vranovensis* or *E. coli*. F2 descendants of P-trained parents showed lower relative preference for *P. vranovensis* than descendants of E-trained parents in *C. elegans* (C) and in *C. remanei* females (B). Centre line in each box: median; box range: 25th to 75th percentiles; whiskers: minimum-maximum values; dots: raw data from choice-assay plates. Grey boxes and dots represent E-trained worms; blue boxes and dots represent P-trained worms. Choice index = (# of worms on *E. coli* – # of worms on *P. vranovensis*)/(# of worms on *E. coli* + # of worms on *P. vranovensis*). n indicates the number of plates included for each species/sex. Positive values indicate preference for *E. coli* (E preference; upward grey arrow) and negative values indicate preference for *P. vranovensis* (P preference; downward blue arrow). The horizontal line at zero indicates no preference.

**Table 4:**
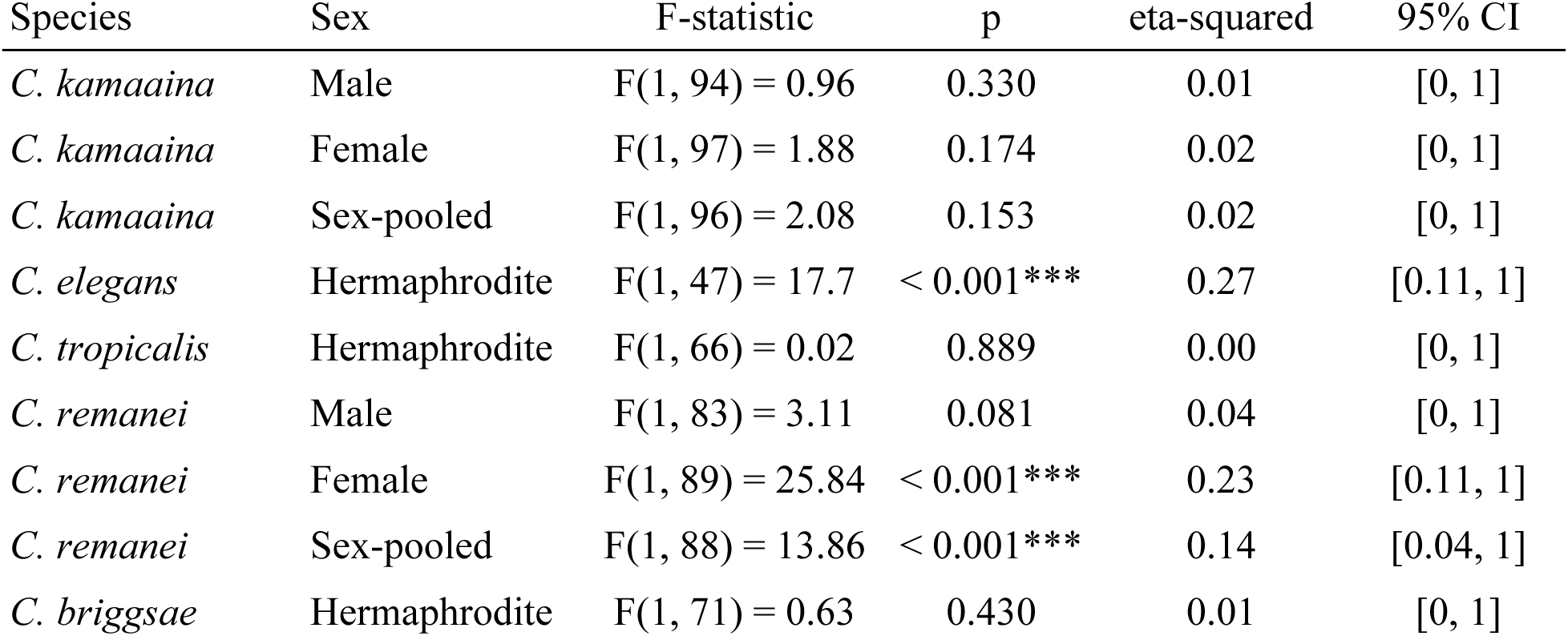
ANOVA statistics, effect sizes (eta-squared) and 95% confidence intervals (CI) for transgenerational response (F2) to parental bacterial training on *P. vranovensis* or *E. coli* across species and sexes.

| Species | Sex | F-statistic | p | eta-squared | 95% CI |
| --- | --- | --- | --- | --- | --- |
| *C. kamaaina* | Male | F(1, 94) = 0.96 | 0.330 | 0.01 | [0, 1] |
| *C. kamaaina* | Female | F(1, 97) = 1.88 | 0.174 | 0.02 | [0, 1] |
| *C. kamaaina* | Sex-pooled | F(1, 96) = 2.08 | 0.153 | 0.02 | [0, 1] |
| *C. elegans* | Hermaphrodite | F(1, 47) = 17.7 | < 0.001*** | 0.27 | [0.11, 1] |
| *C. tropicalis* | Hermaphrodite | F(1, 66) = 0.02 | 0.889 | 0.00 | [0, 1] |
| *C. remanei* | Male | F(1, 83) = 3.11 | 0.081 | 0.04 | [0, 1] |
| *C. remanei* | Female | F(1, 89) = 25.84 | < 0.001*** | 0.23 | [0.11, 1] |
| *C. remanei* | Sex-pooled | F(1, 88) = 13.86 | < 0.001*** | 0.14 | [0.04, 1] |
| *C. briggsae* | Hermaphrodite | F(1, 71) = 0.63 | 0.430 | 0.01 | [0, 1] |

In *C. remanei*, female descendants of P-trained parents showed lower relative preference for *P. vranovensis* than female descendants of E-trained parents, with significant effects and large effect sizes across statistical approaches, whereas no significant difference was detected in males (Figure 6; Table 4; Table S4; see also Supplemental Text and Figure S5). Although E-trained P0 females had significantly higher choice indices than E-trained F2 females, possibly due to differences in experimental handling between generations (Table S5 and Supplemental Text), the contrasts were consistent across experimental blocks (Figure S4 and Figure S5).

In *C. kamaaina*, P-trained P0 males showed a modest increase in relative preference for *E. coli* compared with E-trained P0 males, whereas in F2 neither males nor females differed between descendants of P-trained and E-trained parents (Figure 6; Table 4; Table S4; see also Supplemental Text and Figure S7).

### Transgenerational survival benefits in *C. elegans*, *C. remanei* and *C. tropicalis*

Results of the transgenerational behavioural assays suggested that parental exposure to *P. vranovensis* likely increased descendant fitness through pathogen avoidance. We next asked whether parental exposure to *P. vranovensis* conferred fitness benefit in the descendants when behavioural avoidance was not possible. We therefore measured survival of F2 larvae hatched directly on *P. vranovensis*.

F2 descendants of P-trained worms showed significantly improved survival compared with descendants of E-trained worms in *C. elegans* (Figure 7; Table 5), with a large effect size, as well as in *C. remanei* (Figure 7; Table 5), with a moderate effect size. Notably, in *C. tropicalis*, parental training on *P. vranovensis* also significantly improved survival of F2 descendants on *P. vranovensis* (Figure 7; Table 5), with a large effect size, even though no behavioural change was observed either within or across generations for this species. No survival differences were found in *C. kamaaina* or *C. briggsae* (Figure 7; Table 5).

**Figure 7:**
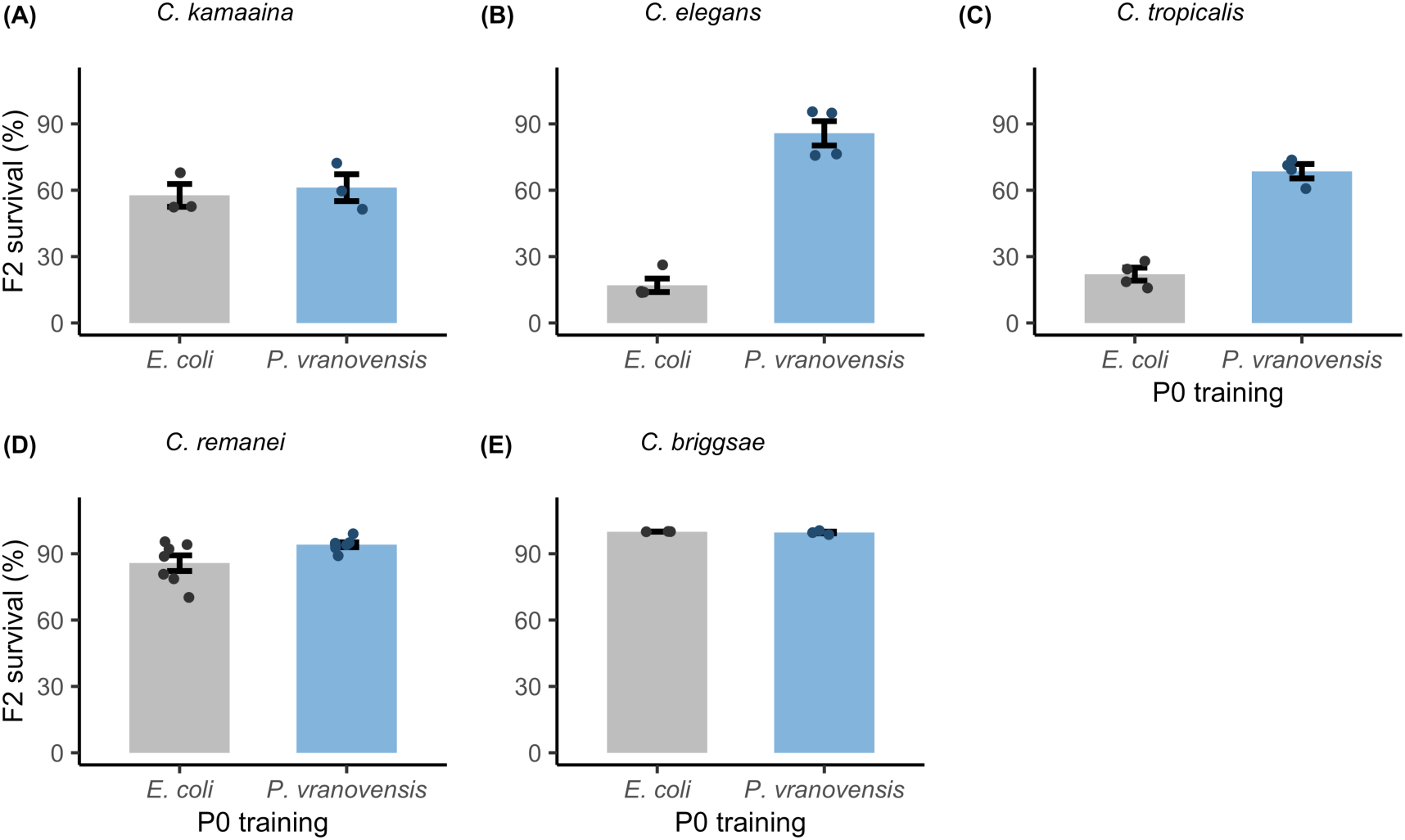
Percentage of newly hatched F2 larvae surviving on *P. vranovensis* in *C. kamaaina* (A), *C. elegans* (B), *C. tropicalis* (C), *C. remanei* (D), and *C. briggsae* (E). Parental training on *P. vranovensis* (blue) significantly increased F2 survival in *C. elegans*, *C. tropicalis*, and *C. remanei* compared with F2 descendants of *E. coli*-trained worms (grey; see Table 5 for more details). Bars and lines: mean values ± standard errors; points: individual plates; n = 3-6 plates per replicate, each with > 80 larvae.

**Table 5:**
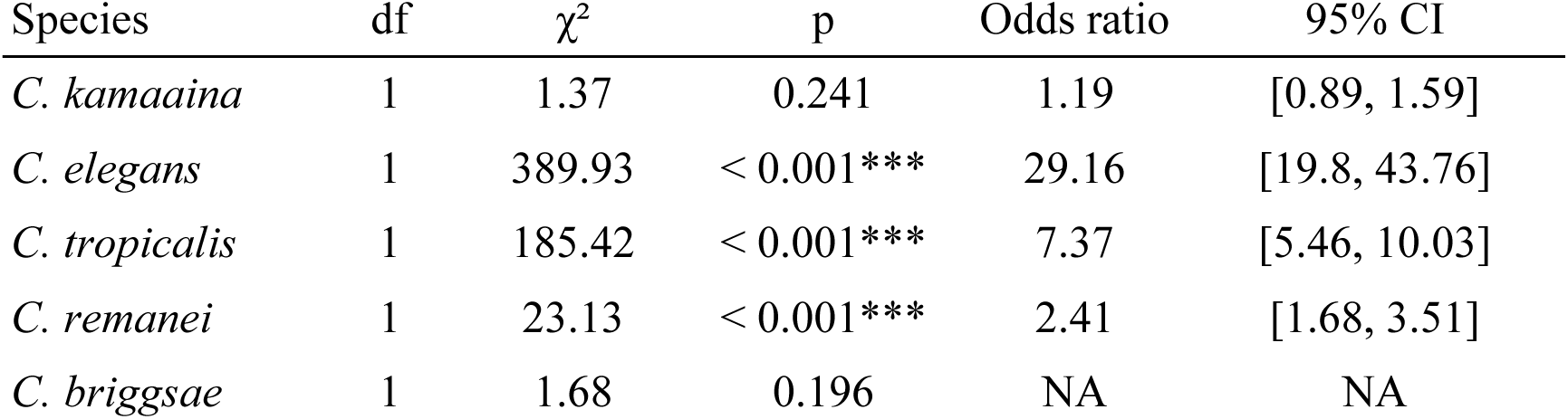
The effect of parental bacterial training (E-versus P-trained parents) on F2 survival on *P. vranovensis* for each of the five species. Odds ratios and 95% CIs are reported as NA for *C. briggsae* because survival was near-complete in both groups, so the estimate was not identifiable.

## Discussion

We showed that behavioural avoidance to *P. vranovensis* pathogen was robustly reproducible in *C. elegans* and did not require azide (Figure 4; Table 2). We then found that this pathogen avoidance varied across *Caenorhabditis* species (Figure 5; Table 3). Moreover, parental pathogen exposure induced adaptive transgenerational effects on offspring pathogen avoidance (Figure 6; Table 4) and survival benefit during subsequent pathogen exposure (Figure 7; Table 5). Further investigation of candidate components of the *C. elegans* TEI pathway suggested that these species-specific responses involve distinct underlying mechanisms (Figure 3).

### Ecological context may shape transgenerational responses to pathogens

Theory predicts that transgenerational responses are favoured when parental exposure reliably predicts descendant exposure^5,6,34,35^. *Caenorhabditis* worms thus are expected to evolve transgenerational responses to pathogens that they naturally and predictably encounter in their habitats. Nonetheless, although *Pseudomonas* bacteria appear to be common in the natural habitats of *C. elegans*^11^, *Caenorhabditis* species differ in their geographical distributions and habitats^36^ and may therefore encounter different microbial communities. Whether and how frequently non-*C. elegans* species encounter *Pseudomonas* bacteria in their natural habitats remain largely unknown.

One possibility is that, in the *Caenorhabditis* species in which we did not observe a transgenerational response to *P. vranovensis*, selection is insufficiently strong or consistent to favour the evolution of such a response. That is, a detrimental but rarely encountered pathogen may provide little opportunity for such a response to evolve. This also implies that transgenerational responses should be specific to particular pathogen species and strains. Indeed, in *C. elegans*, exposure reduced attraction to *P. aeruginosa* PA14, whereas no such response was detected with PAK1 or PAO1 strains^37^. Parental exposure to *P. vranovensis* BIGb0446 increased F1 survival in *C. elegans* and *C. kamaaina* but reduced it in *C. briggsae*^38^. Future studies should therefore establish what pathogens worms encounter in nature, how predictable the exposure to these pathogens is, and whether worms evolve transgenerational responses to these pathogens.

### Different mechanisms may mediate different transgenerational responses

Across species, transgenerational pathogen avoidance was most closely associated with sequence homology between Pv1 and *maco-1,* which may facilitate the downregulation of *maco*-1 and thereby induce behavioural change. Avoidance in F2 was detected in *C. elegans* and *C. remanei*, the two species with perfect or near-perfect homology, but not in *C. kamaaina, C. tropicalis*, and *C. briggsae,* for which the closest match had three or more mismatches (Figure 3). Sensitivity to ingested RNA did not explain within-or transgenerational behavioural responses to pathogen exposure: *C. remanei* showed attraction in P0 and avoidance in F2 despite being insensitive to ingested RNA, whereas *C. kamaaina* was sensitive but showed no detectable response in either P0 or F2^27^. Indeed, previous studies suggest that sensitivity to ingested RNA is not strictly required for behavioural responses to intact bacterial lawns. In *C. elegans*, *sid-2*, *sid-1* and *dcr-1* mediate the uptake, transport and processing of ingested dsRNA, respectively. All three are required for avoidance induced by purified bacterial small RNAs^8,9,13^, but *sid-2* and *dcr-1* mutants can still learn avoidance after exposure to intact bacterial lawns^13^.

Transgenerational survival benefits may be mediated by *cer-1* retrotransposon, which conveys small RNA-based signals across generations. Parental exposure to *P. vranovensis* increased survival in F2 in *C. elegans*, *C. remanei* and *C. tropicalis*, all of which showed evidence of a putatively active *cer-1* retrotransposon (Figure 3 and Figure 7). Overall, our findings suggest that parental exposure may affect pathogen avoidance and survival through different molecular pathways. Direct functional tests will be required to identify the components involved in each response.

### Sex-specificity in within- and transgenerational pathogen responses

Among the two gonochoristic species in our study, *C. remanei* showed within-generation pathogen attraction and transgenerational pathogen avoidance in females but not males (Figure 5 and Figure 6). In *C. kamaaina*, males showed only a modest within-generation response and no detectable transgenerational avoidance, while no behavioural changes were detected in females (Figure 5 and Figure 6). This could be because males, females and hermaphrodites can differ in their responses to microbial challenge. In *C. elegans*, for example, some immune responses are stronger in males, whereas hermaphrodites can rely more on protective microbes under pathogen challenge^39,40^.

Notably, sex-specific expression of a transgenerational phenotype does not necessarily imply sex-specific transmission. Transgenerational effects can be transmitted through both male and female germline in *C. elegans*, yet our results showed that the expression is likely female- and hermaphrodite-specific.

Sex-specificity in transmission and expression may have different consequences in hermaphroditic and gonochoristic species. *Caenorhabditis* species often colonise new resource patches from a small number of founders^26^. In self-fertilising species, an effect transmitted preferentially through one germline can remain within the same lineage^41^, whereas in gonochoristic species its persistence depends on inheritance by the sex capable of transmitting it in successive generations. Mating system may therefore influence how readily transgenerational responses evolve and persist.

### Future work

Our results reveal substantial variation in transgenerational responses even under controlled conditions. Broader comparisons across *Caenorhabditis* lineages, including hermaphroditic and gonochoristic species from the Japonica group, would help establish how these responses evolve. Examining multiple traits across generations and varying rearing and assay conditions may also clarify the sources of this variation. Functional studies are needed to identify the underlying mechanisms. Ultimately, understanding the adaptive significance of these responses requires testing their effects on fitness under ecologically relevant pathogen exposure.

## Conclusion

Our results demonstrate that adaptive transgenerational effects stemming from pathogen exposure can extend beyond inheritance of learned avoidance. We replicated transgenerational pathogen avoidance in *C. elegans* and found that pathogen exposure increased avoidance in F2 female descendants in *C. remanei*. Interestingly, these two species have the closest sequence matches between the bacterial small RNA Pv1 and neuronal target *maco*-1. Crucially, ancestral exposure improved descendant survival in *C. tropicalis* despite the absence of behavioural avoidance in this species. These findings show that ancestral pathogen exposure has both species- and trait-specific transgenerational effects. Therefore, assessing the adaptive value of transgenerational plasticity requires measuring a whole suite of life-history and behavioural traits in descendants. Importantly, the absence of inherited avoidance does not imply the absence of beneficial transgenerational effects.

## Resource availability

### Lead contact

Requests for resources should be directed to the lead contacts, Hwei-yen Chen and Martyna K. Zwoinska, who will fulfil the request.

### Materials availability

No unique reagents were generated in this study.

### Data and code availability

Additional information required to reanalyse the data reported in this paper is available from the lead contact upon request.

## Acknowledgements

We thank members of the *Caenorhabditis* Genomes Project for prepublication access to genome and transcriptome data. We thank the *Caenorhabditis* Genetics Center (CGC) for providing nematode strains and C. T. Murphy at Princeton University for sharing the Pv1 small RNA sequence. *E. coli* OP50 was obtained from the CGC. *P. vranovensis* GRb0427 was a gift from A. A. Maklakov at the University of East Anglia (originally from C. T. Murphy). Figures 1 and 3 were created using BioRender. This work was supported by the Crafoord Foundation to H.Y.C. H.Y.C. was also supported by a Starting Grant from the Swedish Research Council (grant no. 2022-04211). M.I.L. acknowledges funding from the Swedish Research Council (grant no. 2020-04388). M.Z. was supported by the Birgitta Sintring Foundation (S2024-0007) and the Carl Tryggers Foundation (CTS 23:2672).

## Author contributions

Conceptualization: H.Y.C., M.Z., and M.I.L. Methodology: H.Y.C. and M.Z. Investigation: A.D.A., A.S.A., D.A., D.C., E.M.L.D., I.H., N.T.R., A.G.V., and A.N.W. Writing (original draft): H.Y.C., M.Z., and M.I.L. Supervision: H.Y.C., M.Z., and A.A.M. Funding acquisition: H.Y.C.

## Experimental model and study participant details

### Worm maintenance

The following worm species were used: *C. kamaaina* (QG122), *C. elegans* (N2), *C. tropicalis* (JU1373), *C. remanei* (PB4641) and *C. briggsae* (AF16). For all species, stock worms were maintained on 90-mm plates containing nematode growth medium (NGM; 3 g/L NaCl, 2.5 g/L Bacto-peptone, 17 g/L Bacto-agar in distilled water, with the addition of 1 mL/L cholesterol (5 mg/mL in ethanol), 1 mL/L 1 M CaCl2, 1 mL/L 1 M MgSO4, and 25 mL/L 1 M potassium phosphate buffer (pH 6.0) after autoclaving) or high growth medium (HGM; 3 g/L NaCl, 20 g/L Bacto-peptone, 30 g/L Bacto-agar in distilled water, with the addition of 4 mL/L cholesterol (5 mg/mL in ethanol), 1 mL/L 1 M CaCl2, 1 mL/L 1 M MgSO4 and 25 mL/L 1 M KPO4 buffer (pH 6.0) after autoclaving) seeded with 1 mL of *E. coli* OP50 at 20°C throughout the study.

### Bacterial culture

*E. coli* OP50 and *Pseudomonas vranovensis* GRb0427 were used in this study. To prepare liquid bacterial cultures, a 15-mL Falcon tube containing 6 mL of sterile LB broth (10 g/L tryptone + 5 g/L yeast extract + 10 g/L NaCl in distilled water) was inoculated with a single bacterial colony picked from streak plates. Bacterial cultures were then grown at 37°C with shaking at 220 rpm.

## Method details

### Fecundity assay

25 μL of *E. coli* OP50 (OD₆₀₀ = 0.5) was seeded onto 35-mm NGM plates to form a single bacterial lawn in the center of each plate (Figure 1). Day-1 adult hermaphrodites or females with visible mating plugs were washed in M9 buffer and then individually transferred to each plate.

Worms were allowed to lay eggs for 24 hours and were then transferred daily to fresh, seeded plates for three consecutive days. After the mothers were removed each day, plates were incubated for 48 hours to allow the offspring to hatch and develop into larvae. Plates were then heated at 85°C for 15 minutes to kill the offspring, and age-specific fecundity was measured as the number of larvae produced each day.

### Survival assay

1 mL of an overnight culture of *P. vranovensis* GRb0427 was seeded onto 60-mm NGM plates with the bacterial lawn covering the entire surface. Nematode eggs were bleached onto seeded plates and incubated for 24 hours at 20°C. After incubation, the numbers of live and dead worms outside the bleaching spot were counted to calculate percentage survival.

#### Bacterial training

Bacterial training and aversive learning assays followed published protocols^9,29^ with minor modifications (Figure 1). To prepare bacterial training plates, overnight bacterial cultures of *E. coli* and *P. vranovensis* were standardised in sterile LB broth to an optical density (OD₆₀₀) of 0.5, and 800 μL (UEA) or 1 mL (UU, LU) of a standardised bacterial culture was seeded onto 90-mm NGM plates with the bacterial lawn covering the entire surface. Plates were dried under a sterile hood and then incubated at 25°C for 48 hours before being used.

To prepare P0 worms, healthy populations (i.e., no contamination or crowding, and at least one week after thawing) were bleached to derive synchronised eggs. Eggs were plated onto NGM or HGM plates seeded with *E. coli* OP50 and allowed to develop to the L4 larval stage (ca. 48-52 hours post-synchronisation).

On the day of training, all training plates were equilibrated to room temperature (ca. 20°C) for about 1 hour before the addition of worms. Synchronised L4 worms were washed twice and transferred to training plates. Specifically, worms were washed off plates using ca. 5 mL of M9 buffer (6 g/L Na2HPO4, 3 g/L KH2PO4, 5 g/L NaCl and 1 mL/L 1 M MgSO4 in distilled water) into a 15-mL Falcon tube and allowed to settle by gravity (< 5 minutes). Once the worms had settled, the supernatant was discarded, and 5 mL of fresh M9 buffer was added to each tube to remove residual bacteria from their body surfaces. Total washing time was kept under 30 minutes to minimise the risk of stressing or starving the worms.

After removing the supernatant from the second wash, a small amount of M9 buffer (ca. 200-500 μL) was added to resuspend the worm pellet. Worms were then pipetted onto the training plates. For *E. coli* training, a maximum of 800 worms were transferred to each training plate to avoid crowding or running out of food. For *P. vranovensis* training, ca. 1600 worms were added to each training plate to account for mortality. Worms were incubated on training plates for 24 hours at 20°C. All training plates were right-side up, but plates from different training treatments were stored in separate containers.

### Aversive learning assay

To prepare bacterial choice-assay plates (see Figure 1), 25 μL of a standardised bacterial culture (OD₆₀₀ = 0.5) of *E. coli* or *P. vranovensis* was spotted onto opposite sides of a 60-mm NGM plate, with approximately 2 cm between the two spots. The left/right position of each bacterial species was alternated between plates. Plates were dried under a sterile hood and then incubated at 25°C for 48 hours prior to use.

On the day of the assay, plates were equilibrated to room temperature for about 1 hour before the addition of worms. Worms were washed as described in Bacterial training. However, the supernatant from the first wash was retained for collection of the next generation.

For aversive learning assays testing the effect of potassium azide, 1 µL of 1 M potassium azide was pipetted onto each bacterial spot immediately before the assay. In all other assays, no potassium azide was added. Washed worms were transferred in M9 buffer to the bottom centre of the choice plate, midway between and equidistant from the two bacterial spots. Excess M9 buffer was carefully removed using a Kimwipe to release the worms. Plates were then incubated undisturbed in the dark for 1 hour.

After incubation, plates were heated at 85°C for 15 minutes to kill the worms and prevent further movement. For assays using potassium azide, we counted all worms within 2 mm of each bacterial lawn. For assays without azide, we counted worms on each bacterial lawn as well as those within 1 mm of its edge. Choice index was calculated for each replicate as (# of worms on *E. coli* − # of worms on *P. vranovensis*)/(total # of worms scored on the two bacterial lawns).

Plates containing fewer than 20 worms were excluded for sex-pooled analyses in male–female species and hermaphrodites, while plates with fewer than 10 individuals of the focal sex were excluded for sex-specific analyses in male–female species. When necessary, images of bacterial spots were captured with a camera mounted on a Nikon SMZ1270 microscope using NIS-Elements software, and worms were counted from the images using the open-source FIJI software.

### Transgenerational behavioural assay

To test behavioural changes in F2 descendants, the retained wash supernatant was bleached to obtain F1 embryos (Figure 1). F1 animals were allowed to develop for 3 days at 20°C until early adulthood, after which they were bleached to obtain F2 embryos. F2 worms were grown for 3 days until day-1 adulthood and were then subjected to the choice assay as described above, including choice-index calculation and quantification. Both P0 and F2 animals were tested as day-1 adults. For statistical analysis, P0 and F2 data were each pooled across two to three independent experimental blocks.

### Pv1-maco-1 homology analysis

We aimed to identify *Caenorhabditis* species whose *maco*-1 ortholog contained the previously reported 16-nt sequence homology between the bacterial Pv1 small RNA and the *C. elegans maco*-1 gene (sequence name: D2092.5). We focused on species within the *Elegans* group. We retrieved protein and CDS sequences from all species available on WormBase^42^ (release WS293^49^) and the *Caenorhabditis* Genomes Project^43^ (v2^50^). Species and datasets used are listed in Table S7. Orthogroups of protein sequences from all species were constructed using OrthoFinder^44^ (v3.0.1b1^51^, default options). SeqKit^45^ (v2.8.0^52^, seqkit locate –max-mismatch 3) was used to identify the 16-nt Pv1 sequence among the CDSs of all sequences belonging to the orthogroup containing MACO-1, allowing up to three nucleotide mismatches. For each match, we documented the number of mismatches, the matched sequence, its position within the ortholog, and the strand on which it occurred.

### Sequence analysis of cer-1 retrotransposons

To check for the presence of *cer*-1 LTR retrotransposons in *C. tropicalis, C. remanei*, *C. briggsae* and *C. kamaaina*, reference assemblies were obtained from WormBase^42^ (*C. tropicalis*, *C. briggsae*) and Zenodo (*C. kamaaina*; Zenodo record 12633738). For all species, *de novo* repeat libraries were built with RepeatModeler^46^ (2.0.7^53^), using the -LTRStruct flag to enable LTR-specific structural detection. Genomes were then soft-masked with RepeatMasker^47^.

Candidate LTRs were identified structurally using LTRharvest (GenomeTools^48^ 1.6.6^55^) and protein domains were annotated with LTRdigest using the GyDB HMM database^49^ and the WormBase *C. elegans* tRNA database to detect primer-binding sites. Additionally, *C. elegans* Cer-1 protein sequences (GAG aa 1–734, POL aa 735–1774, CERV aa 1775–2272) were used as tBLASTn queries against each reference genome. Candidate *cer*-1 sequences were classified as high-confidence (POL homology >35% identity over ≥100 aa, intact LTR structure, and PBS motif TGGGGGCCG with up to two mismatches), partial (lacking one or two hallmarks), or no evidence. All LTRdigest candidates were screened against the *C. elegans* proteome to remove false positives caused by host genes.

### Statistical analysis

All analyses were conducted in R version 4.5.2 (2025-10-31) (R Core Team), using the packages lme4 (1.1.38), glmmTMB (1.1.14), rstatix (0.7.3), effectsize (1.0.2), and car (3.1.5). Plots were generated using ggplot2 (4.0.2).

#### Fecundity

**λ**_ind_ was analysed using independent two-sample t-tests comparing bacterial training conditions. For each test, we extracted the t-statistic, degrees of freedom and p-value. Effect sizes were quantified using Hedges’ g, along with corresponding 95% confidence intervals. Age-specific fecundity was analysed using generalised linear mixed-effects models (GLMMs) with a negative binomial error distribution to account for overdispersion in count data. Total offspring production was modelled as a function of bacterial training condition, age (centred), and age squared, including their interactions. Random intercepts for plate were included to account for non-independence among individuals within plates. Model significance was assessed using Type II Wald chi-square tests using the car package.

#### Choice index

Choice index (CI) was assessed using both parametric and non-parametric methods. Specifically, we performed one-way ANOVAs, two-sided Student’s t-tests and Wilcoxon rank-sum tests to evaluate differences between training conditions. Only plates meeting the worm-count criteria specified in the Aversive learning assay section were included. Effect sizes were quantified as eta-squared for ANOVA, Hedges’ g for t-tests and rank-biserial correlation coefficients (r) for Wilcoxon tests.

To better account for the discrete and bounded nature of the data, we additionally modelled behavioural responses as binomial outcomes using generalised linear models (GLMs) or generalised linear mixed-effects models (GLMMs). These models used the counts of worms on each bacterial lawn (*E. coli* or *P. vranovensis*) as the response, rather than the derived CI. When multiple experimental blocks were present, we fitted GLMMs with a binomial error distribution and logit link function, including training condition as a fixed effect and block as a random intercept. For datasets containing a single block, we instead fitted binomial GLMs without random effects, with training condition as the fixed effect. Statistical significance of model terms in GLMs and GLMMs was assessed using Type II Wald chi-square tests using the car package. Effect sizes for GLM and GLMM analyses are reported as odds ratios with 95% confidence intervals obtained from model estimates. Across all analyses, reported p-values are two-sided.

#### Survival

Survival was analysed using generalised linear models (GLMs) with a binomial error distribution and logit link function. Survival counts (numbers of live versus dead individuals) were modelled as a function of bacterial training, and significance was assessed using Type II Wald chi-square tests using the car package. Effect sizes were quantified as odds ratios with 95% confidence intervals obtained from the fitted models. Extreme estimates (e.g., due to complete absence of mortality in one group) were not interpreted.

## Supplemental Materials

### Supplemental Figures and Tables

**Figure S1:**
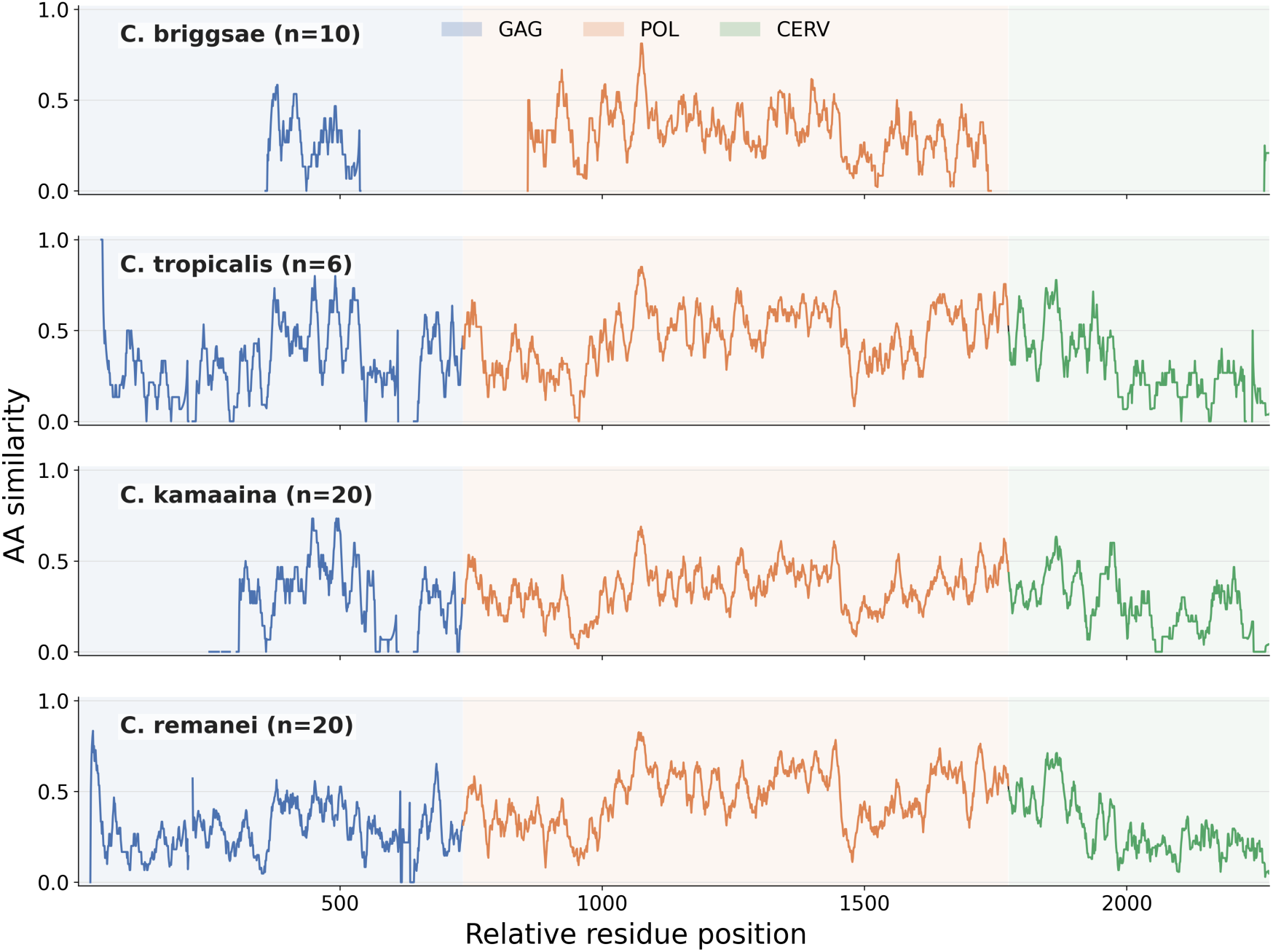
Protein sequence similarities of the GAG, POL, and CERV regions among *C. briggsae*, *C. tropicalis*, *C. kamaaina*, and *C. remanei*.

**Table S1:**
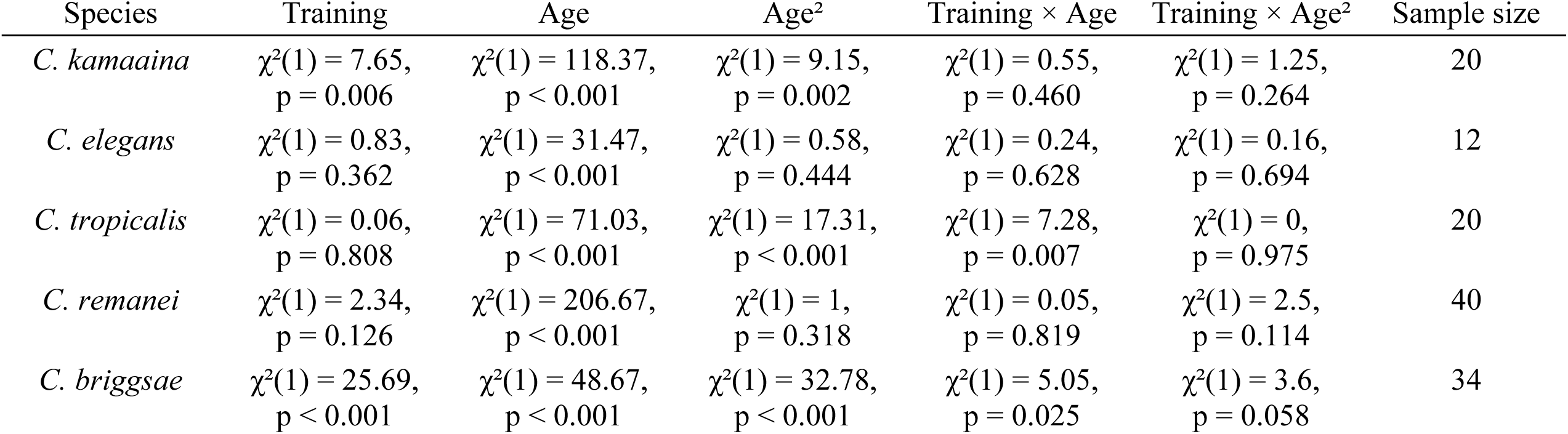
Type II Wald chi-square tests from generalised linear mixed-effects models evaluating the effects of bacterial training, age, and age^2^ on age-specific fecundity in the P0 generation across all five *Caenorhabditis* species. Models included bacterial training, age, age^2^, and their interactions as fixed effects, and plate as a random effect. Sample size indicates the number of plates included for each species.

| Species | Training | Age | Age<sup>2</sup> | Training × Age | Training × Age<sup>2</sup> | Sample size |
| --- | --- | --- | --- | --- | --- | --- |
| *C. kamaaina* | $\chi^2(1) = 7.65$,<br>p = 0.006 | $\chi^2(1) = 118.37$,<br>p < 0.001 | $\chi^2(1) = 9.15$,<br>p = 0.002 | $\chi^2(1) = 0.55$,<br>p = 0.460 | $\chi^2(1) = 1.25$,<br>p = 0.264 | 20 |
| *C. elegans* | $\chi^2(1) = 0.83$,<br>p = 0.362 | $\chi^2(1) = 31.47$,<br>p < 0.001 | $\chi^2(1) = 0.58$,<br>p = 0.444 | $\chi^2(1) = 0.24$,<br>p = 0.628 | $\chi^2(1) = 0.16$,<br>p = 0.694 | 12 |
| *C. tropicalis* | $\chi^2(1) = 0.06$,<br>p = 0.808 | $\chi^2(1) = 71.03$,<br>p < 0.001 | $\chi^2(1) = 17.31$,<br>p < 0.001 | $\chi^2(1) = 7.28$,<br>p = 0.007 | $\chi^2(1) = 0$,<br>p = 0.975 | 20 |
| *C. remanei* | $\chi^2(1) = 2.34$,<br>p = 0.126 | $\chi^2(1) = 206.67$,<br>p < 0.001 | $\chi^2(1) = 1$,<br>p = 0.318 | $\chi^2(1) = 0.05$,<br>p = 0.819 | $\chi^2(1) = 2.5$,<br>p = 0.114 | 40 |
| *C. briggsae* | $\chi^2(1) = 25.69$,<br>p < 0.001 | $\chi^2(1) = 48.67$,<br>p < 0.001 | $\chi^2(1) = 32.78$,<br>p < 0.001 | $\chi^2(1) = 5.05$,<br>p = 0.025 | $\chi^2(1) = 3.6$,<br>p = 0.058 | 34 |

**Table S2:**
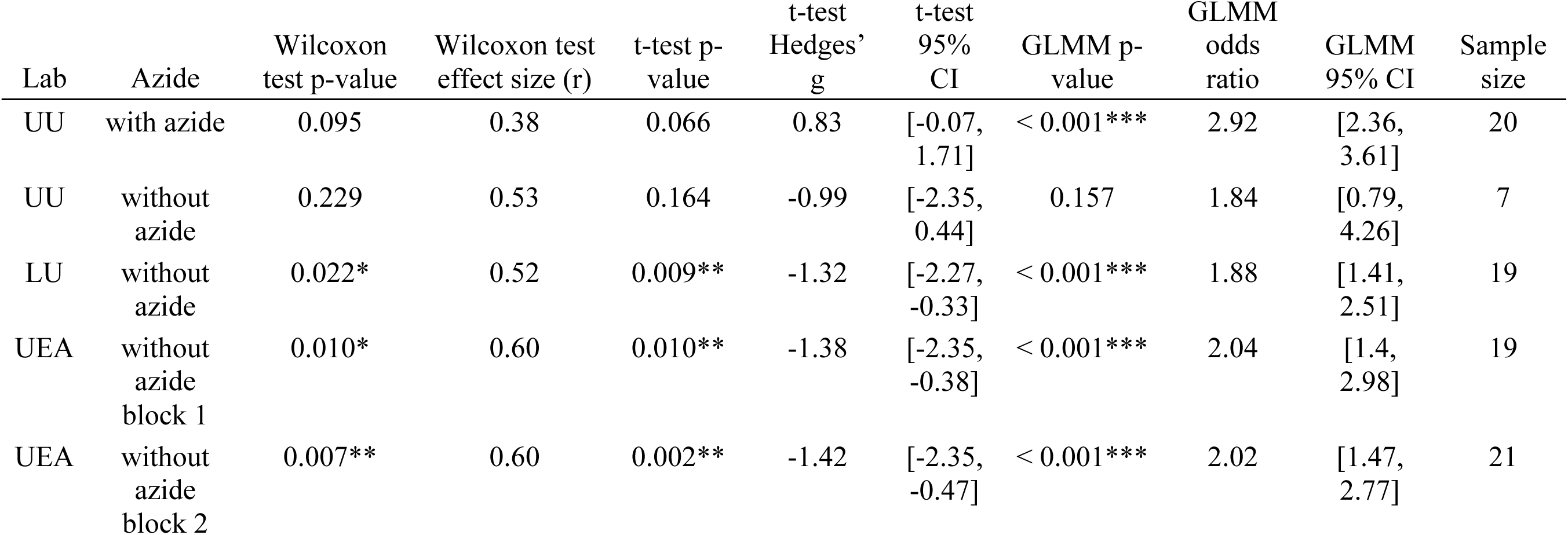
Parental pathogen avoidance in *C. elegans* with or without azide, analysed by Wilcoxon rank-sum tests, t-tests, and generalised linear models. The parental learned-avoidance response was assessed independently at three laboratories: Uppsala University (UU) and Lund University (LU), Sweden, and the University of East Anglia (UEA), UK. Sample size indicates the number of plates included in each dataset.

| Lab | Azide | Wilcoxon<br>test p-value | Wilcoxon test<br>effect size (r) | t-test p-value | t-test<br>Hedges' g | t-test<br>95%<br>CI | GLMM p-value | GLMM<br>odds<br>ratio | GLMM<br>95% CI | Sample<br>size |
| --- | --- | --- | --- | --- | --- | --- | --- | --- | --- | --- |
| UU | with azide | 0.095 | 0.38 | 0.066 | 0.83 | [-0.07,<br>1.71] | < 0.001*** | 2.92 | [2.36,<br>3.61] | 20 |
| UU | without<br>azide | 0.229 | 0.53 | 0.164 | -0.99 | [-2.35,<br>0.44] | 0.157 | 1.84 | [0.79,<br>4.26] | 7 |
| LU | without<br>azide | 0.022* | 0.52 | 0.009** | -1.32 | [-2.27,<br>-0.33] | < 0.001*** | 1.88 | [1.41,<br>2.51] | 19 |
| UEA | without<br>azide<br>block 1 | 0.010* | 0.60 | 0.010** | -1.38 | [-2.35,<br>-0.38] | < 0.001*** | 2.04 | [1.4,<br>2.98] | 19 |
| UEA | without<br>azide<br>block 2 | 0.007** | 0.60 | 0.002** | -1.42 | [-2.35,<br>-0.47] | < 0.001*** | 2.02 | [1.47,<br>2.77] | 21 |

**Figure S2:**
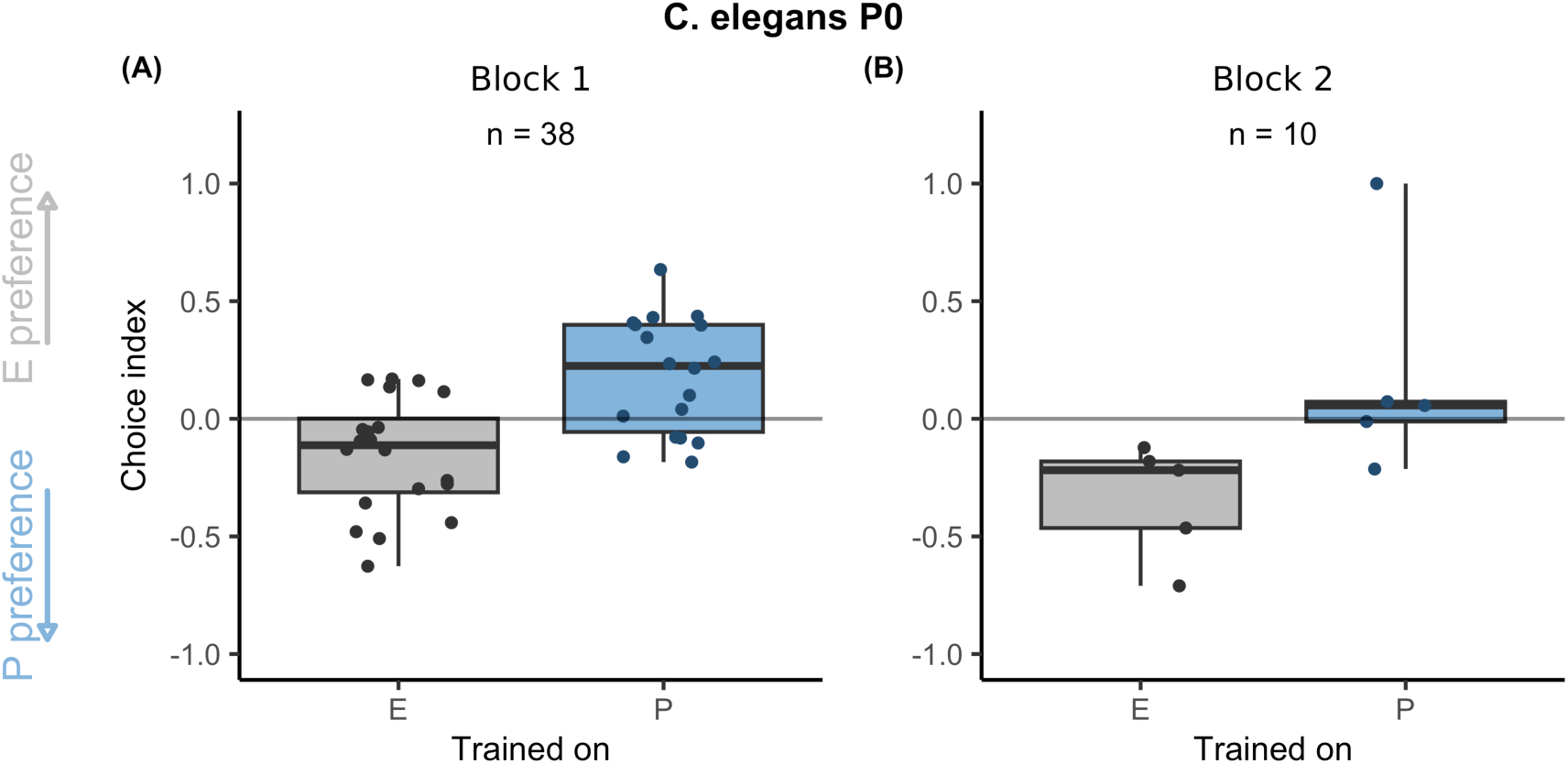
Behavioural responses of E-trained and P-trained worms of *C. elegans* in the P0 generation across experimental blocks. Assays were performed without potassium azide. Centre line in each box: median; box range: 25th to 75th percentiles; whiskers: minimum-maximum values; dots: raw data from choice-assay plates. Grey boxes and dots represent E-trained worms; blue boxes and dots represent P-trained worms. n indicates the number of plates included in each dataset. Choice index = (# of worms on *E. coli* – # of worms on *P. vranovensis*)/(# of worms on *E. coli* + # of worms on *P. vranovensis*). Positive values indicate preference for *E. coli* (E preference; upward grey arrow) and negative values indicate preference for *P. vranovensis* (P preference; downward blue arrow). The horizontal line at zero indicates no preference.

**Figure S3:**
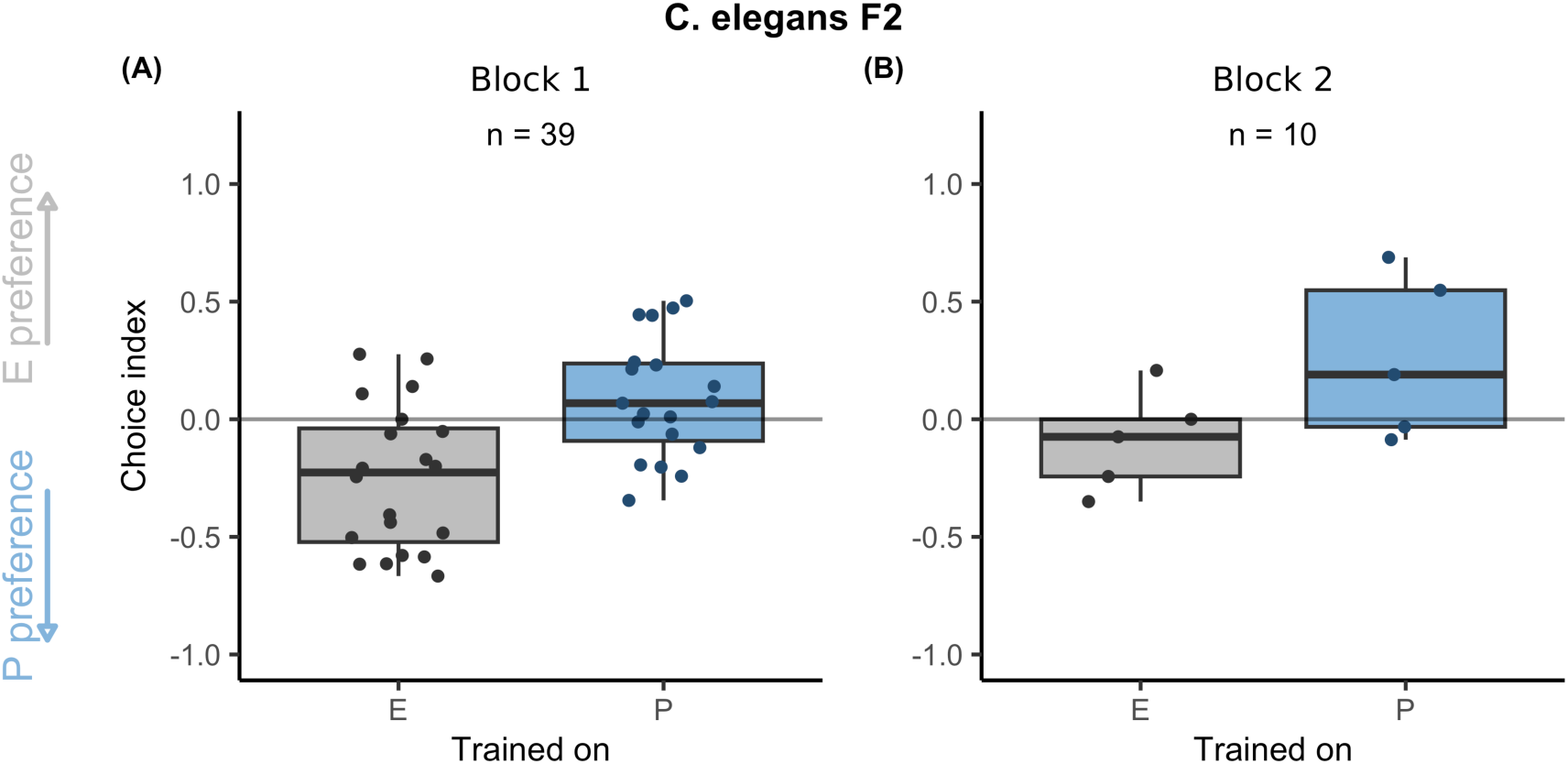
Behavioural responses of E-trained and P-trained worms of *C. elegans* in the F2 generation across experimental blocks. Assays were performed without potassium azide. Centre line in each box: median; box range: 25th to 75th percentiles; whiskers: minimum-maximum values; dots: raw data from choice-assay plates. Grey boxes and dots represent E-trained worms; blue boxes and dots represent P-trained worms. n indicates the number of plates included in each dataset. Choice index = (# of worms on *E. coli* – # of worms on *P. vranovensis*)/(# of worms on *E. coli* + # of worms on *P. vranovensis*). Positive values indicate preference for *E. coli* (E preference; upward grey arrow) and negative values indicate preference for *P. vranovensis* (P preference; downward blue arrow). The horizontal line at zero indicates no preference.

**Figure S4:**
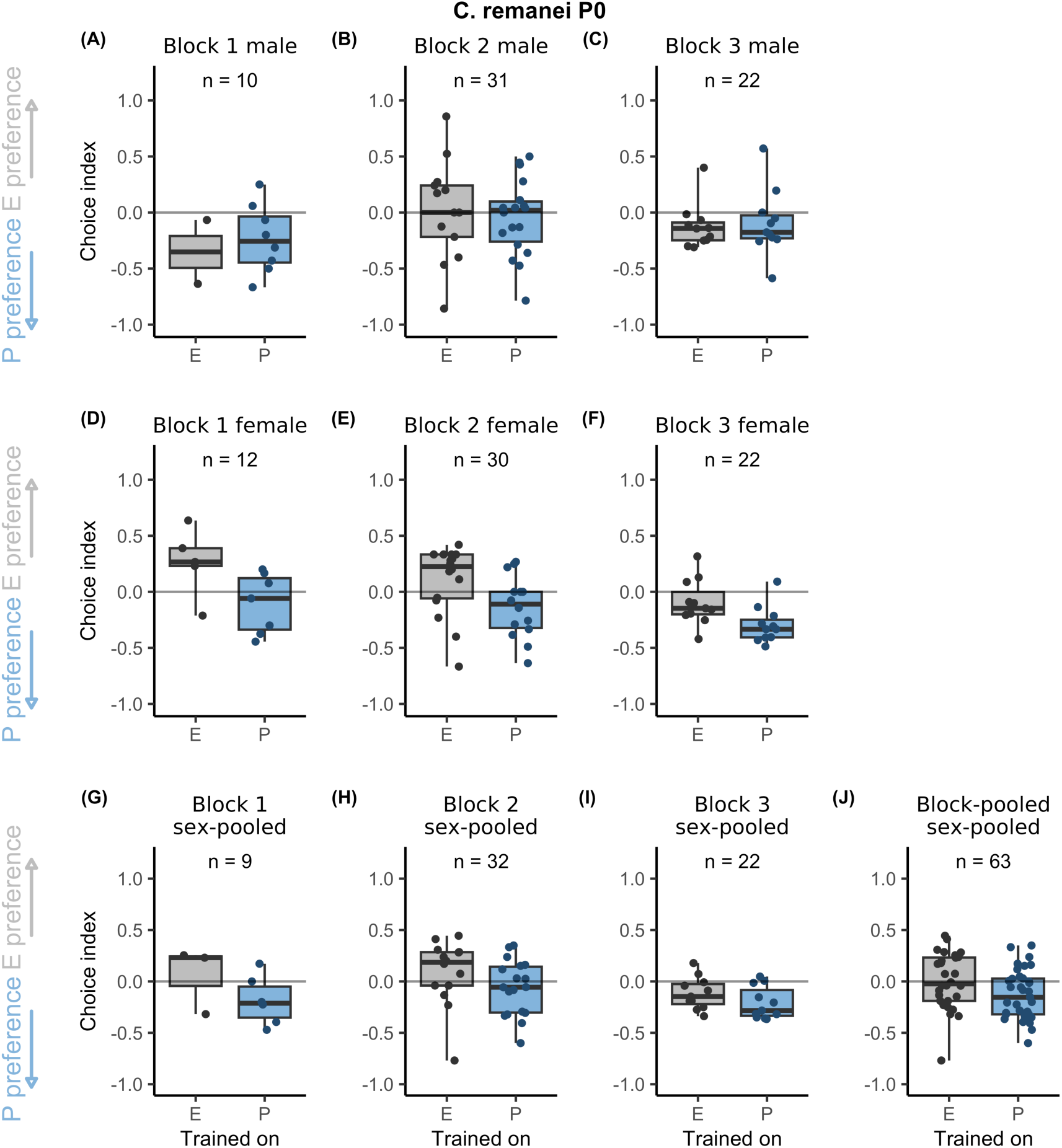
Behavioural responses of E-trained and P-trained *C. remanei* males and females in the P0 generation across experimental blocks.s Assays were performed without potassium

**Figure S5:**
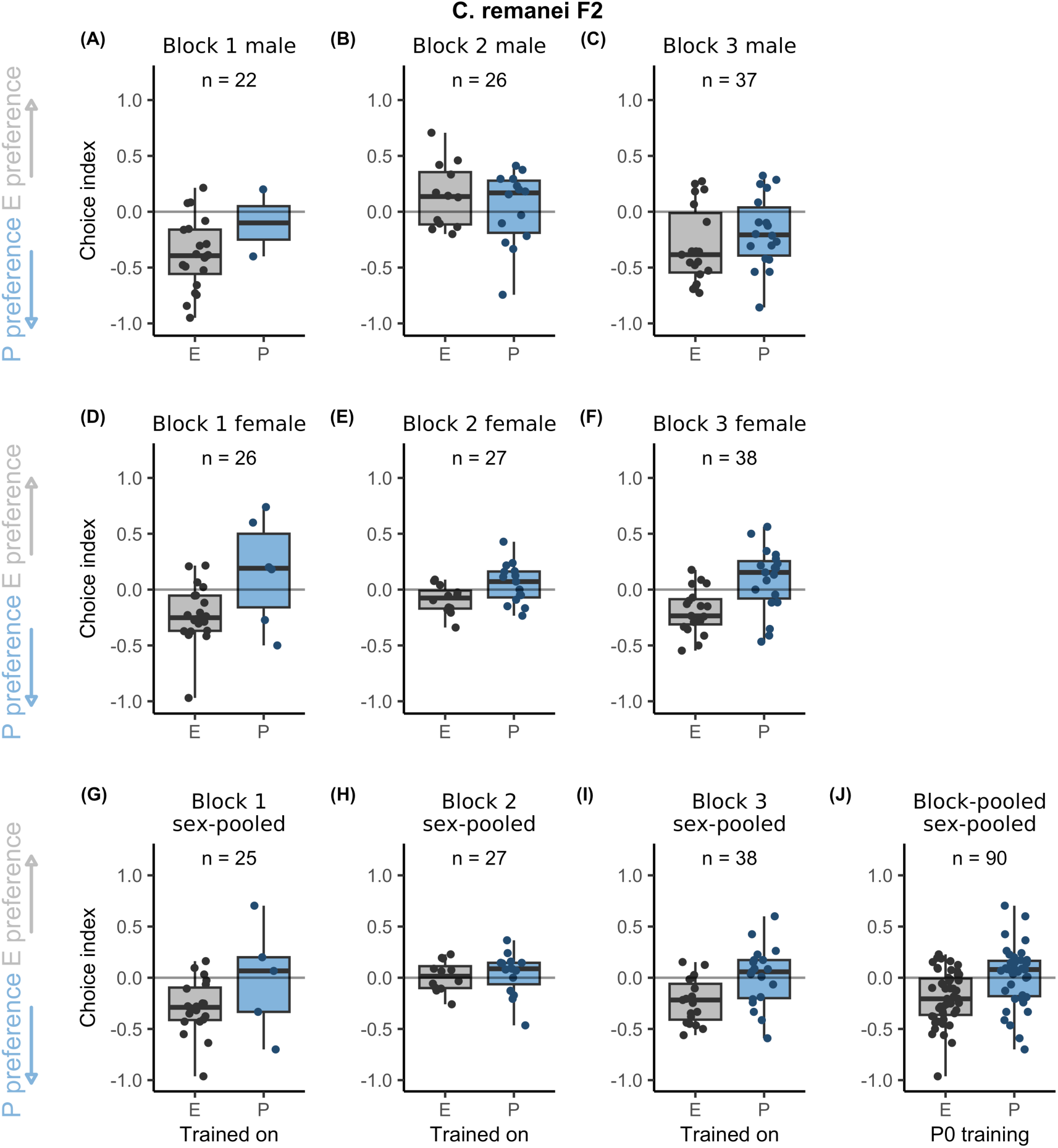
Behavioural responses of E-trained and P-trained *C. remanei* males and females in the F2 generation across experimental blocks. Assays were performed without potassium

**Figure S6:**
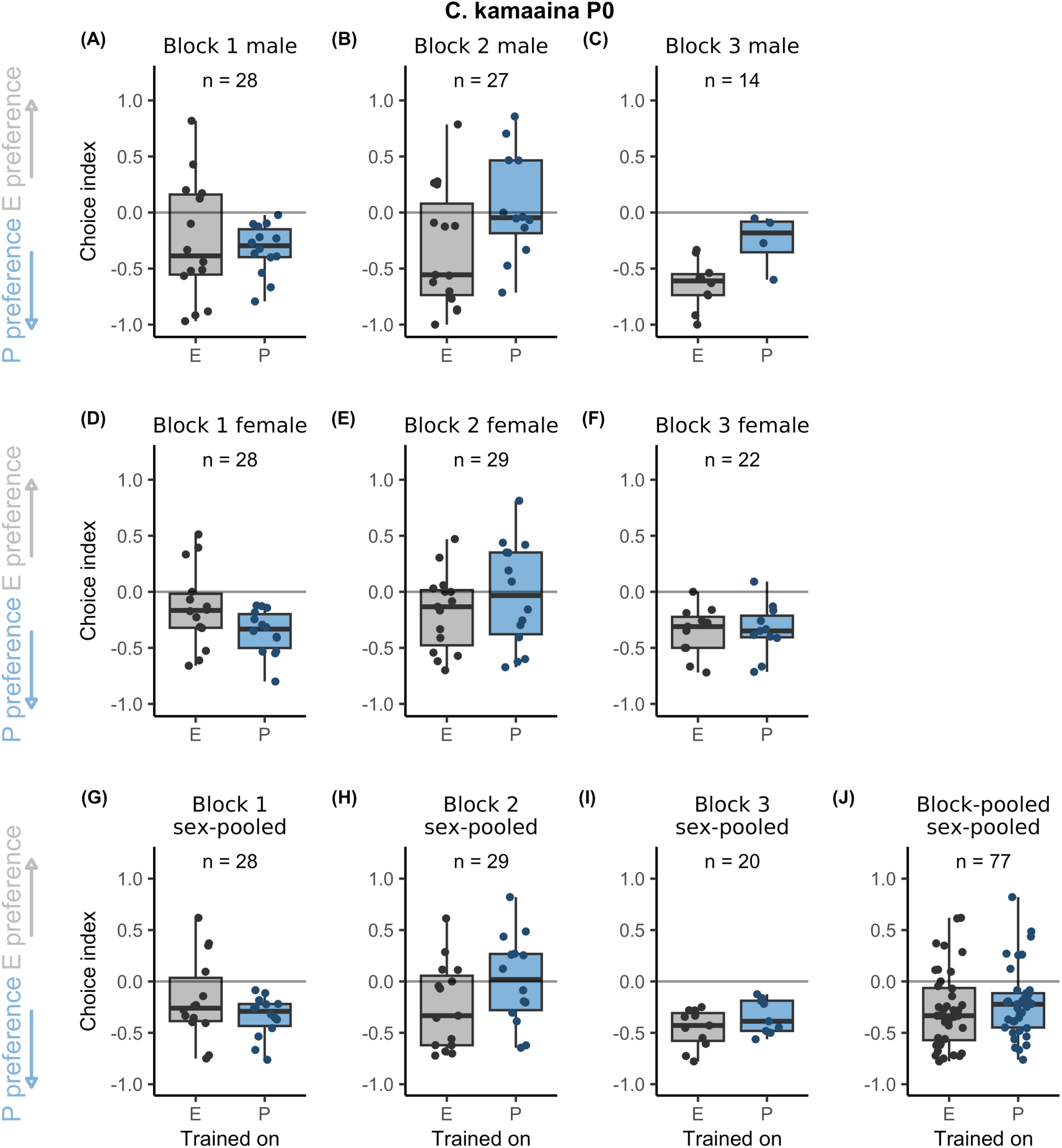
Behavioural responses of E-trained and P-trained *C. kamaaina* males and females in the P0 generation across experimental blocks. Assays were performed without potassium

**Figure S7:**
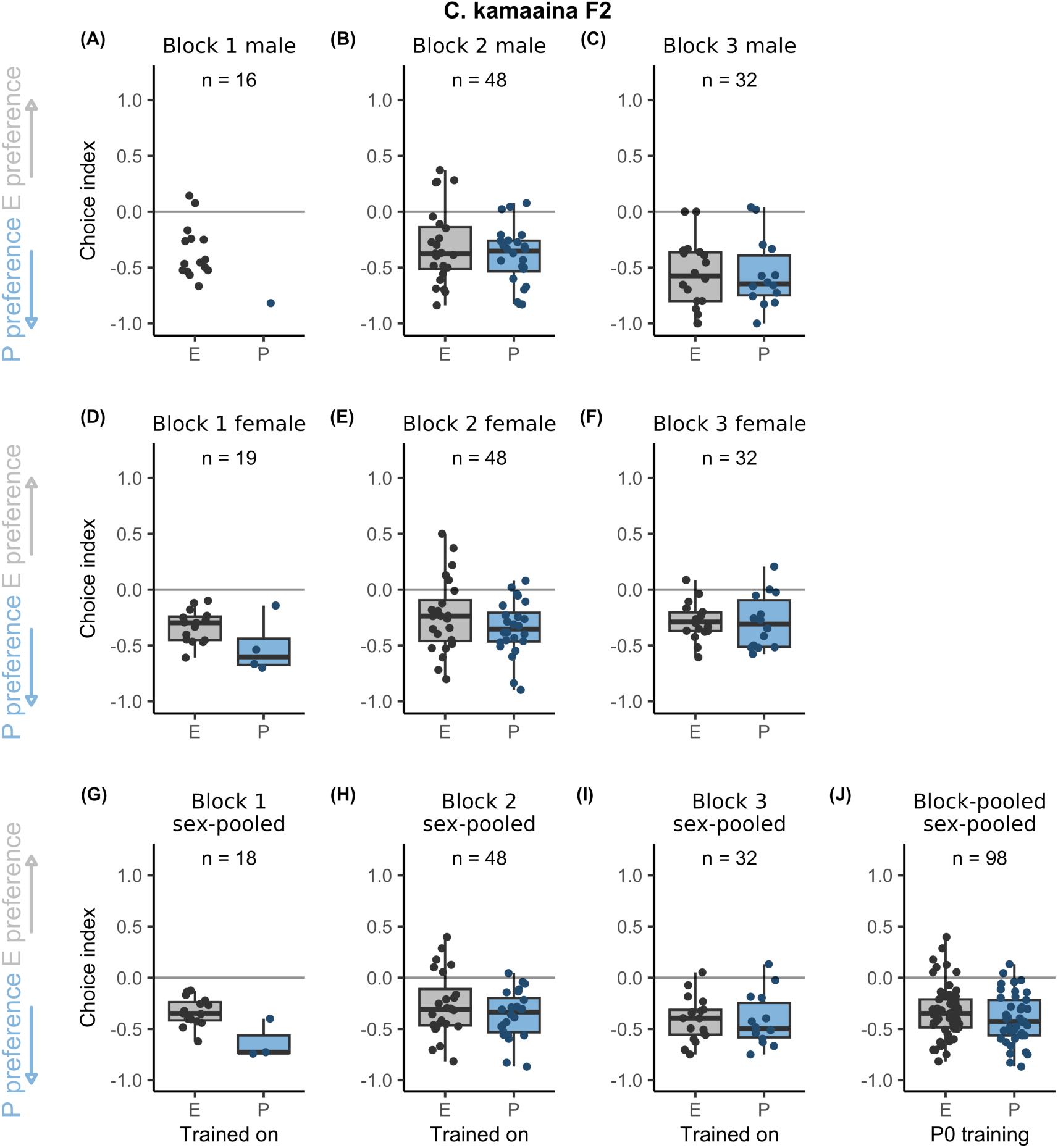
Behavioural responses of E-trained and P-trained *C. kamaaina* males and females in the F2 generation across experimental blocks. Assays were performed without potassium

**Table S3:**
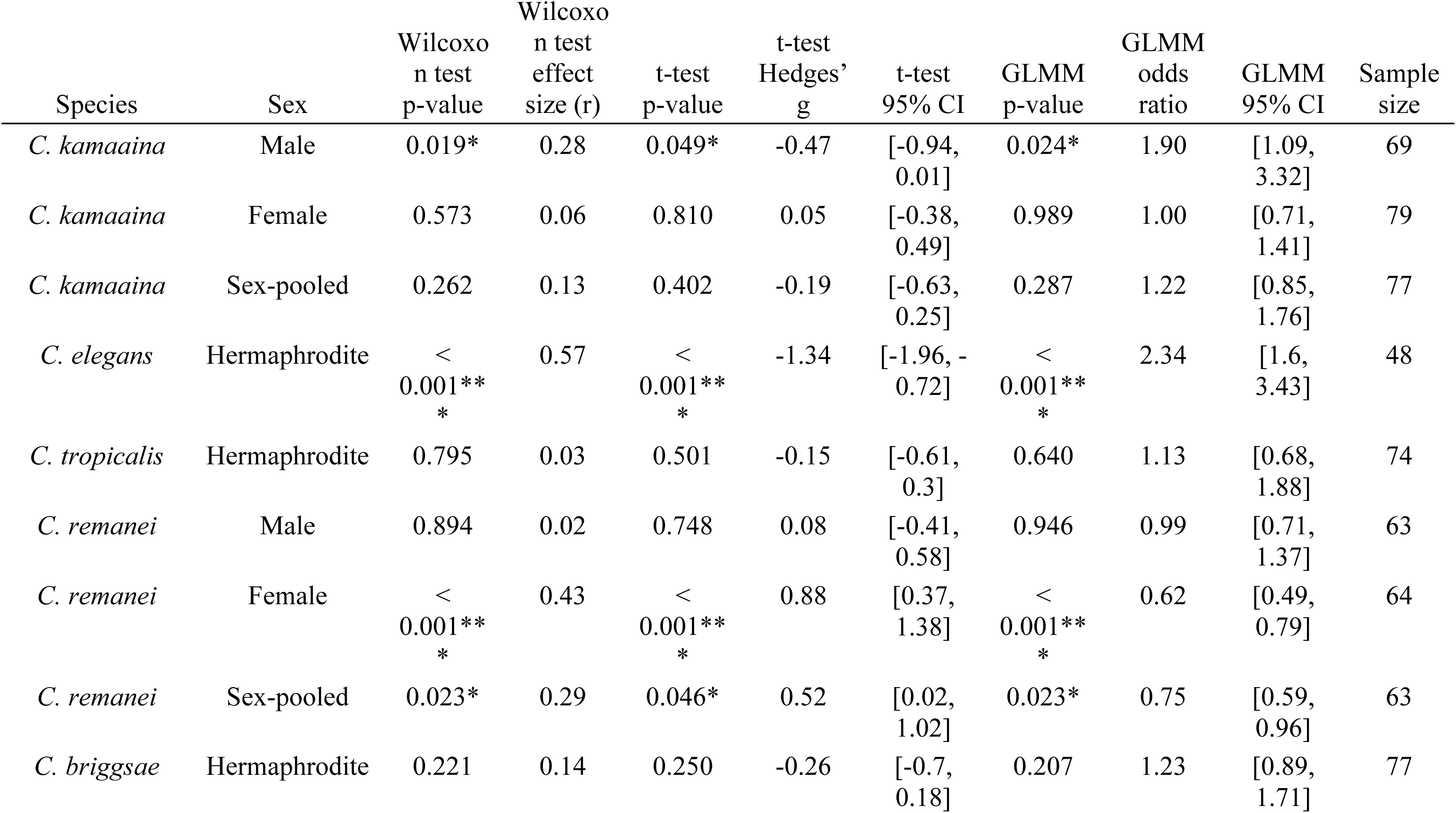
Differences in behavioural responses between E-trained and P-trained worms in each nematode species in the P0 generation, analysed by Wilcoxon rank-sum tests, t-tests, and generalised linear mixed-effects models (with bacterial training fitted as a fixed effect and plate nested within experimental block included as a random effect). Sample size indicates the number of plates included for each species.

**Table S4:**
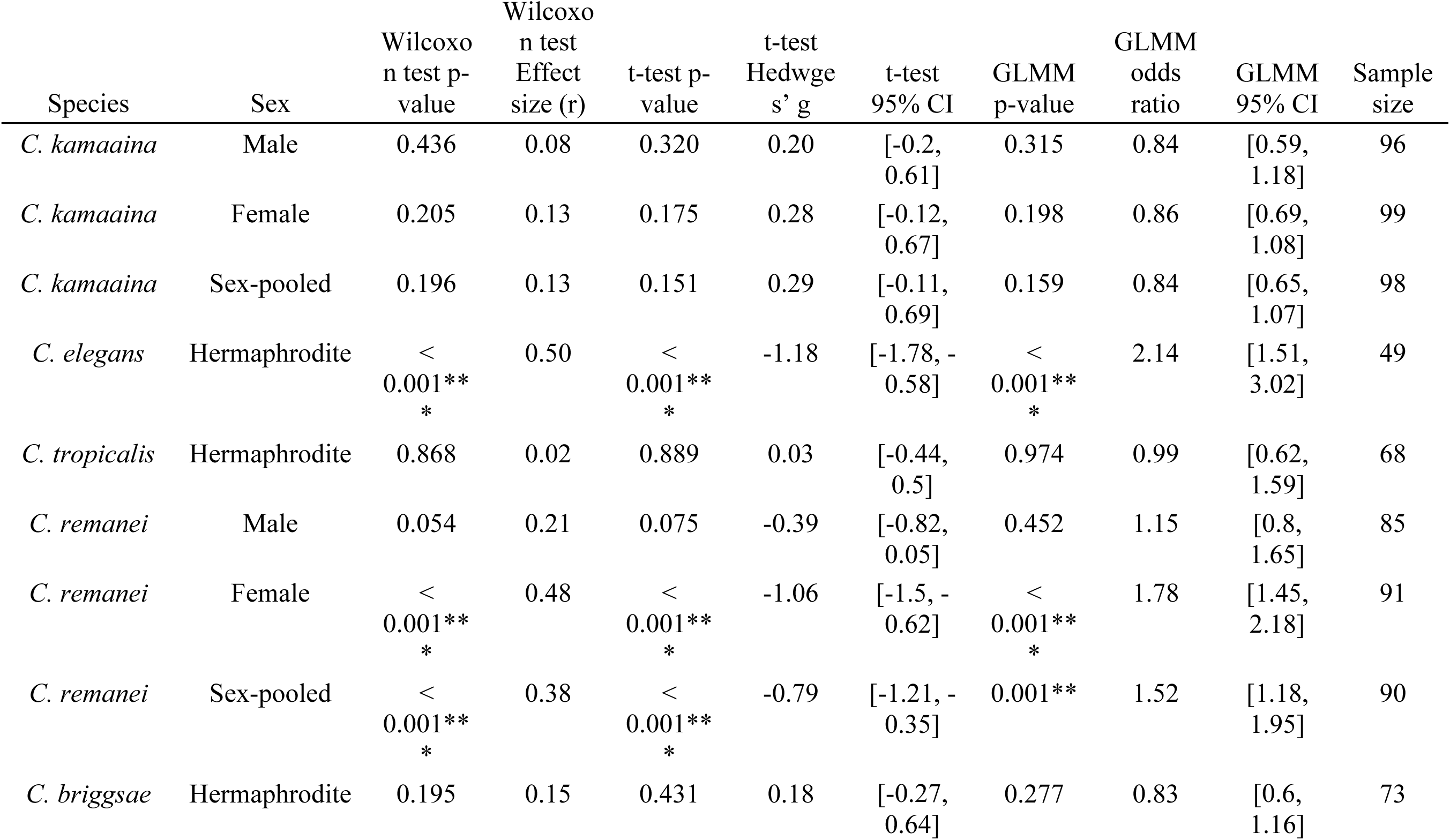
Effect of parental bacterial training on F2 behavioural responses in each nematode species, analysed by Wilcoxon rank-sum tests, t-tests, and generalised linear mixed-effects models (with bacterial training fitted as a fixed effect and plate nested within experimental block included as a random effect). Sample size indicates the number of plates included for each species.

**Table S5:** ANOVA statistics, effect sizes (η^2^), and 95% confidence intervals (CI) for differences in choice index between P0 and F2 generations in E-trained animals across species and sexes.

| Species | Sex | F-statistic | p | eta-squared | 95% CI |
| --- | --- | --- | --- | --- | --- |
| *C. kamaaina* | Male | F(1, 94) = 0.11 | 0.743 | 0.00 | [0, 1] |
| *C. kamaaina* | Female | F(1, 95) = 1.06 | 0.307 | 0.01 | [0, 1] |
| *C. elegans* | Hermaphrodite | F(1, 48) = 0.14 | 0.710 | 0.00 | [0, 1] |
| *C. tropicalis* | Hermaphrodite | F(1, 69) = 13.47 | < 0.001*** | 0.16 | [0.05, 1] |
| *C. remanei* | Male | F(1, 75) = 3.06 | 0.084 | 0.04 | [0, 1] |
| *C. remanei* | Female | F(1, 81) = 18.18 | < 0.001*** | 0.18 | [0.07, 1] |
| *C. briggsae* | Hermaphrodite | F(1, 55) = 3.33 | 0.073 | 0.06 | [0, 1] |

**Table S6:**
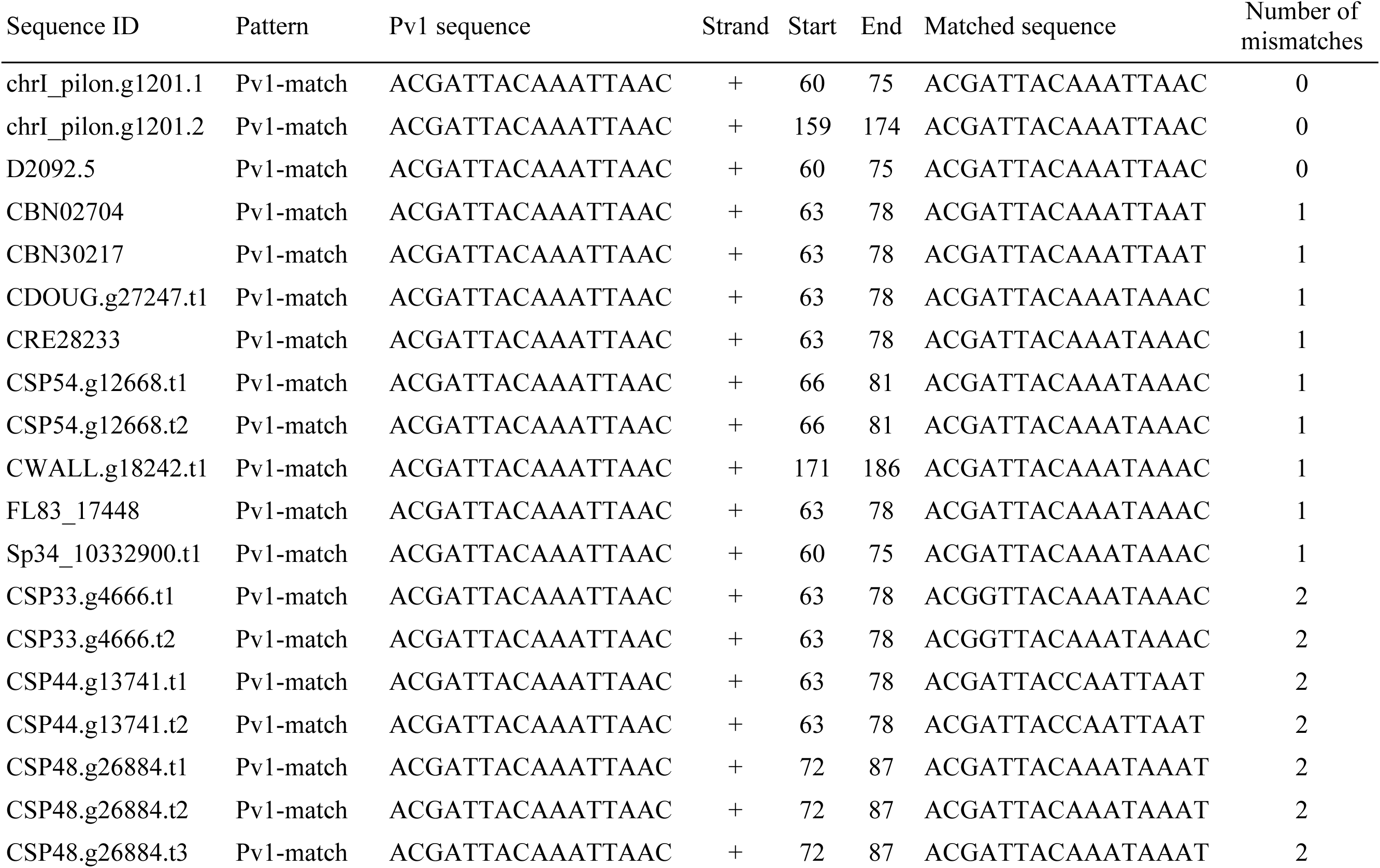
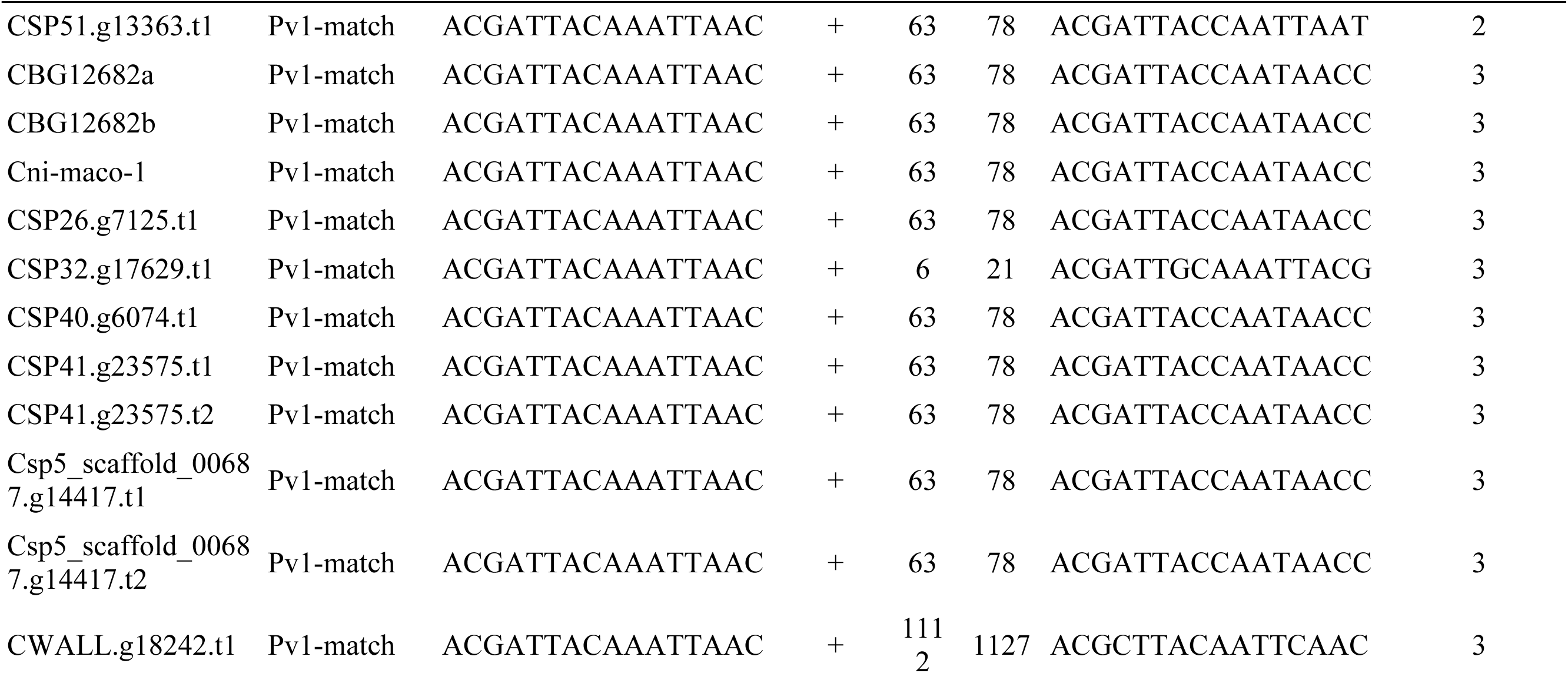
Sequence matches to the Pv1 sequence in *Caenorhabditis* nematodes.

**Table S7:** *Caenorhabditis* genomic resources used for Pv1 sequence searches.

| Source | Release | Species | NCBI accession | CDS file | Protein file |
| --- | --- | --- | --- | --- | --- |
| Caenorhabditis Genomes Project | v2 | *Caenorhabditis afra* | NA | CAFRA.caenorhabditis_afra_JU1286_v2.cd<br>s.fna | CAFRA.caenorhabditis_afra_JU1286_v2.proteins.faa |
| Caenorhabditis Genomes Project | v2 | *Caenorhabditis astrocarya* | NA | CASTR.caenorhabditis_astrocarya_NIC1040_v2.cds.fna | CASTR.caenorhabditis_astrocarya_NIC1040_v2.proteins.faa |
| Caenorhabditis Genomes Project | v2 | *Caenorhabditis auriculariae* | NA | CAURI.caenorhabditis_auriculariae_NKZ352_PRJEB40642.cds.fna | CAURI.caenorhabditis_auriculariae_NKZ352_PRJEB40642.proteins.faa |
| Caenorhabditis Genomes Project | v2 | *Caenorhabditis castelli* | NA | CCAST.caenorhabditis_castelli_JU1956_v2.cds.fna | CCAST.caenorhabditis_castelli_JU1956_v2.proteins.faa |
| Caenorhabditis Genomes Project | v2 | *Caenorhabditis dolens* | NA | CDOLE.caenorhabditis_dolens_NIC394_v2.cds.fna | CDOLE.caenorhabditis_dolens_NIC394_v2.proteins.faa |
| Caenorhabditis Genomes Project | v2 | *Caenorhabditis doughertyi* | NA | CDOUG.caenorhabditis_doughertyi_JU1771_v2.cds.fna | CDOUG.caenorhabditis_doughertyi_JU1771_v2.proteins.faa |
| Caenorhabditis Genomes Project | v2 | *Caenorhabditis drosophilae* | NA | CDROS.caenorhabditis_drosophilae_DF5077_v2.cds.fna | CDROS.caenorhabditis_drosophilae_DF5077_v2.proteins.faa |
| Caenorhabditis Genomes Project | v2 | *Caenorhabditis imperialis* | NA | CIMPE.caenorhabditis_imperialis_EG5942_v2.cds.fna | CIMPE.caenorhabditis_imperialis_EG5942_v2.proteins.faa |
| Caenorhabditis Genomes Project | v2 | *Caenorhabditis kamaaina* | NA | CKAMA.caenorhabditis_kamaaina_QG2077_v2.cds.fna | CKAMA.caenorhabditis_kamaaina_QG2077_v2.proteins.faa |
| Caenorhabditis Genomes Project | v2 | *Caenorhabditis macrosperma* | NA | CMACR.caenorhabditis_macrosperma_JU2083_v2.cds.fna | CMACR.caenorhabditis_macrosperma_JU2083_v2.proteins.faa |
| Caenorhabditis Genomes Project | v2 | *Caenorhabditis monodelphis* | NA | CMONO.caenorhabditis_monodelphis_JU1667_v2.cds.fna | CMONO.caenorhabditis_monodelphis_JU1667_v2.proteins.faa |
| Caenorhabditis Genomes Project | v2 | *Caenorhabditis nouraguensis* | NA | CNOUR.caenorhabditis_nouraguensis_JU2079_v2.cds.fna | CNOUR.caenorhabditis_nouraguensis_JU2079_v2.proteins.faa |
| Caenorhabditis Genomes Project | v2 | *Caenorhabditis oiwi* | NA | COIWI.caenorhabditis_oiwi_ECA1100_v2.cds.fna | COIWI.caenorhabditis_oiwi_ECA1100_v2.proteins.faa |
| Caenorhabditis Genomes Project | v2 | *Caenorhabditis plicata* | NA | CPLIC.caenorhabditis_plicata_SB355_v2.cd<br>s.fna | CPLIC.caenorhabditis_plicata_SB355_v2.proteins.faa |
| Caenorhabditis Genomes Project | v2 | *Caenorhabditis portoensis* | NA | CPORT.caenorhabditis_portoensis_EG5626_v2.cds.fna | CPORT.caenorhabditis_portoensis_EG5626_v2.proteins.faa |
| Caenorhabditis Genomes Project | v2 | *Caenorhabditis* sp2 | NA | CSP02.caenorhabditis_sp2_DF5070_v2.cds.fna | CSP02.caenorhabditis_sp2_DF5070_v2.proteins.faa |
| Caenorhabditis Genomes Project | v2 | *Caenorhabditis* sp8 | NA | CSP08.caenorhabditis_sp8_DF5173_v2.cds.fna | CSP08.caenorhabditis_sp8_DF5173_v2.proteins.faa |
| Caenorhabditis Genomes Project | v2 | *Caenorhabditis* sp24 | NA | CSP24.caenorhabditis_sp24_QG555_v2.cds.fna | CSP24.caenorhabditis_sp24_QG555_v2.proteins.faa |
| Caenorhabditis Genomes Project | v2 | *Caenorhabditis* sp25 | NA | CSP25.caenorhabditis_sp25_ZF1457_v2.cds.fna | CSP25.caenorhabditis_sp25_ZF1457_v2.proteins.faa |
| Caenorhabditis Genomes Project | v2 | *Caenorhabditis* sp27 | NA | CSP27.caenorhabditis_sp27_ECA211_v2.cd<br>s.fna | CSP27.caenorhabditis_sp27_ECA211_v2.proteins.faa |
| Caenorhabditis Genomes Project | v2 | *Caenorhabditis* sp30 | NA | CSP30.caenorhabditis_sp30_DF5174_v2.cds.fna | CSP30.caenorhabditis_sp30_DF5174_v2.proteins.faa |
| Caenorhabditis Genomes Project | v2 | *Caenorhabditis* sp33 | NA | CSP33.caenorhabditis_sp33_BRC20065_v2.cds.fna | CSP33.caenorhabditis_sp33_BRC20065_v2.proteins.faa |
| Caenorhabditis Genomes Project | v2 | *Caenorhabditis* sp41 | NA | CSP41.caenorhabditis_sp41_BRC20276_v2.cds.fna | CSP41.caenorhabditis_sp41_BRC20276_v2.proteins.faa |
| Caenorhabditis Genomes Project | v2 | *Caenorhabditis* sp44 | NA | CSP44.caenorhabditis_sp44_BRC20300_v2.cds.fna | CSP44.caenorhabditis_sp44_BRC20300_v2.proteins.faa |
| Caenorhabditis Genomes Project | v2 | *Caenorhabditis* sp48 | NA | CSP48.caenorhabditis_sp48_BRC20454_v2.cds.fna | CSP48.caenorhabditis_sp48_BRC20454_v2.proteins.faa |
| Caenorhabditis Genomes Project | v2 | *Caenorhabditis* sp49 | NA | CSP49.caenorhabditis_sp49_BRC20456_v2.cds.fna | CSP49.caenorhabditis_sp49_BRC20456_v2.proteins.faa |
| Caenorhabditis Genomes Project | v2 | *Caenorhabditis* sp51 | NA | CSP51.caenorhabditis_sp51_QG2939_v2.cds.fna | CSP51.caenorhabditis_sp51_QG2939_v2.proteins.faa |
| Caenorhabditis Genomes Project | v2 | *Caenorhabditis* sp54 | NA | CSP54.caenorhabditis_sp54_BRC20483_v2.cds.fna | CSP54.caenorhabditis_sp54_BRC20483_v2.proteins.faa |
| Caenorhabditis Genomes Project | v2 | *Caenorhabditis* sp56 | NA | CSP56.caenorhabditis_sp56_JU2215_v2.cds.fna | CSP56.caenorhabditis_sp56_JU2215_v2.proteins.faa |
| Caenorhabditis Genomes Project | v2 | *Caenorhabditis virilis* | NA | CVIRI.caenorhabditis_virilis_JU1968_v2.cds.fna | CVIRI.caenorhabditis_virilis_JU1968_v2.proteins.faa |
| Caenorhabditis Genomes Project | v2 | *Caenorhabditis wallacei* | NA | CWALL.caenorhabditis_wallacei_JU1898_v2.cds.fna | CWALL.caenorhabditis_wallacei_JU1898_v2.proteins.faa |
| Caenorhabditis Genomes Project | v2 | *Caenorhabditis yunquensis* | NA | CYUNQ.caenorhabditis_yunquensis_XZ1518_v2.cds.fna | CYUNQ.caenorhabditis_yunquensis_XZ1518_v2.proteins.faa |
| WormBase | WS293 | *B. malayi* | PRJNA10729 | b_malayi.PRJNA10729.WS293.CDS_transcripts.fa.gz | b_malayi.PRJNA10729.WS293.protein.fa.gz |
| WormBase | WS293 | *C. angaria* | PRJNA51225 | c_angaria.PRJNA51225.WS293.CDS_transcripts.fa.gz | c_angaria.PRJNA51225.WS293.protein.fa.gz |
| WormBase | WS293 | *C. becei* | PRJEB28243 | c_becei.PRJEB28243.WS293.CDS_transcripts.fa.gz | c_becei.PRJEB28243.WS293.protein.fa.gz |
| WormBase | WS293 | *C. bovis* | PRJEB34497 | c_bovis.PRJEB34497.WS293.CDS_transcripts.fa.gz | c_bovis.PRJEB34497.WS293.protein.fa.gz |
| WormBase | WS293 | *C. brenneri* | PRJNA20035 | c_brenneri.PRJNA20035.WS293.CDS_transcripts.fa.gz | c_brenneri.PRJNA20035.WS293.protein.fa.gz |
| WormBase | WS293 | *C. briggsae* | PRJNA10731 | c_briggsae.PRJNA10731.WS293.CDS_transcripts.fa.gz | c_briggsae.PRJNA10731.WS293.protein.fa.gz |
| WormBase | WS293 | *C. elegans* | PRJEB28388 | c_elegans.PRJEB28388.WS293.CDS_transcripts.fa.gz | c_elegans.PRJEB28388.WS293.protein.fa.gz |
| WormBase | WS293 | *C. elegans* | PRJNA13758 | c_elegans.PRJNA13758.WS293.CDS_transcripts.fa.gz | c_elegans.PRJNA13758.WS293.protein.fa.gz |
| WormBase | WS293 | *C. inopinata* | PRJDB5687 | c_inopinata.PRJDB5687.WS293.CDS_transcripts.fa.gz | c_inopinata.PRJDB5687.WS293.protein.fa.gz |
| WormBase | WS293 | *C. japonica* | PRJNA12591 | c_japonica.PRJNA12591.WS293.CDS_transcripts.fa.gz | c_japonica.PRJNA12591.WS293.protein.fa.gz |
| WormBase | WS293 | *C. latens* | PRJNA248912 | c_latens.PRJNA248912.WS293.CDS_transcripts.fa.gz | c_latens.PRJNA248912.WS293.protein.fa.gz |
| WormBase | WS293 | *C. nigoni* | PRJNA384657 | c_nigoni.PRJNA384657.WS293.CDS_transcripts.fa.gz | c_nigoni.PRJNA384657.WS293.protein.fa.gz |
| WormBase | WS293 | *C. panamensis* | PRJEB28259 | c_panamensis.PRJEB28259.WS293.CDS_transcripts.fa.gz | c_panamensis.PRJEB28259.WS293.protein.fa.gz |
| WormBase | WS293 | *C. parvicauda* | PRJEB12595 | c_parvicauda.PRJEB12595.WS293.CDS_transcripts.fa.gz | c_parvicauda.PRJEB12595.WS293.protein.fa.gz |
| WormBase | WS293 | *C. quiockensis* | PRJEB11354 | c_quiockensis.PRJEB11354.WS293.CDS_transcripts.fa.gz | c_quiockensis.PRJEB11354.WS293.protein.fa.gz |
| WormBase | WS293 | *C. remanei* | PRJNA53967 | c_remanei.PRJNA53967.WS293.CDS_transcripts.fa.gz | c_remanei.PRJNA53967.WS293.protein.fa.gz |
| WormBase | WS293 | *C. remanei* | PRJNA577507 | c_remanei.PRJNA577507.WS293.CDS_transcripts.fa.gz | c_remanei.PRJNA577507.WS293.protein.fa.gz |
| WormBase | WS293 | *C. sinica* | PRJNA194557 | c_sinica.PRJNA194557.WS293.CDS_transcripts.fa.gz | c_sinica.PRJNA194557.WS293.protein.fa.gz |
| WormBase | WS293 | *C. sulstoni* | PRJEB12601 | c_sulstoni.PRJEB12601.WS293.CDS_transcripts.fa.gz | c_sulstoni.PRJEB12601.WS293.protein.fa.gz |
| WormBase | WS293 | *C. tribulationis* | PRJEB12608 | c_tribulationis.PRJEB12608.WS293.CDS_transcripts.fa.gz | c_tribulationis.PRJEB12608.WS293.protein.fa.gz |
| WormBase | WS293 | *C. tropicalis* | PRJNA53597 | c_tropicalis.PRJNA53597.WS293.CDS_transcripts.fa.gz | c_tropicalis.PRJNA53597.WS293.protein.fa.gz |
| WormBase | WS293 | *C. uteleia* | PRJEB12600 | c_uteleia.PRJEB12600.WS293.CDS_transcripts.fa.gz | c_uteleia.PRJEB12600.WS293.protein.fa.gz |
| WormBase | WS293 | *C. waitukubuli* | PRJEB12602 | c_waitukubuli.PRJEB12602.WS293.CDS_transcripts.fa.gz | c_waitukubuli.PRJEB12602.WS293.protein.fa.gz |
| WormBase | WS293 | *C. zanzibari* | PRJEB12596 | c_zanzibari.PRJEB12596.WS293.CDS_transcripts.fa.gz | c_zanzibari.PRJEB12596.WS293.protein.fa.gz |
| WormBase | WS293 | *O. tipulae* | PRJEB15512 | o_tipulae.PRJEB15512.WS293.CDS_transcripts.fa.gz | o_tipulae.PRJEB15512.WS293.protein.fa.gz |
| WormBase | WS293 | *O. volvulus* | PRJEB513 | o_volvulus.PRJEB513.WS293.CDS_transcripts.fa.gz | o_volvulus.PRJEB513.WS293.protein.fa.gz |
| WormBase | WS293 | *P. pacificus* | PRJNA12644 | p_pacificus.PRJNA12644.WS293.CDS_transcripts.fa.gz | p_pacificus.PRJNA12644.WS293.protein.fa.gz |
| WormBase | WS293 | *P. redivivus* | PRJNA186477 | p_redivivus.PRJNA186477.WS293.CDS_transcripts.fa.gz | p_redivivus.PRJNA186477.WS293.protein.fa.gz |
| WormBase | WS293 | *S. ratti* | PRJEB125 | s_ratti.PRJEB125.WS293.CDS_transcripts.fa.gz | s_ratti.PRJEB125.WS293.protein.fa.gz |
| WormBase | WS293 | *T. muris* | PRJEB126 | t_muris.PRJEB126.WS293.CDS_transcripts.fa.gz | t_muris.PRJEB126.WS293.protein.fa.gz |

## Supplemental Text

### Variability of choice indices across experimental blocks and generations

Variability across experimental blocks is likely due to the experimental protocol’s sensitivity to environmental conditions^14^. Factors such as temperature, light and handling can affect the results (see discussion below). Between-block and between-laboratory variation has also been reported in studies of transgenerational pathogen avoidance^19,20^. Although we followed the protocol as consistently as possible, some variation between blocks was unavoidable. Nonetheless, the magnitude of the variation in our study was comparable to previous reports^9^, and the direction of the effects was consistent across experimental blocks.

Variability was also observed between the P0 and F2 generations, particularly in the choice index of E-trained worms. In *C. remanei* females, the magnitude of the P0 training effect may therefore have been influenced by variation in the E-trained baseline. However, P-trained females showed attraction to *P. vranovensis* in absolute terms, and the direction of the effect of training was consistent across experimental blocks.

We argue that this difference likely reflects variation in pre-assay handling between P0 and F2 generations. Specifically, P0 parents underwent more washing steps than F2 descendants: P0 individuals were transferred from HGM plates to bacterial training plates at the L4 stage, and subsequently from the training plates to choice plates 24 hours later as young adults. In contrast, F2 descendants were not subjected to bacterial training and were transferred directly from HGM plates to choice plates as young adults. Although we made efforts to minimise handling time, washing is known to be a stressful procedure for worms^29^, and the additional washing steps in P0 may have contributed to the observed differences.

We believe that both within- and transgenerational behavioural effects observed in our study reflect genuine biological responses for the following reasons. First, the variation we observed between P0 and F2 generations is similar to patterns reported in previous studies using *P. aeruginosa*^10^, suggesting that such variability is a general feature of this experimental paradigm rather than an artefact of our specific setup. Second, E- and P-trained worms were compared within each generation, where both groups underwent the same procedures. This means that the observed differences are unlikely to arise from experimental handling. Finally, the differences between E-trained and P-trained worms were consistently observed across independent experimental blocks. Together, these observations support the reproducibility of the effect of training while also showing that baseline choice behaviour is sensitive to experimental context.

Factors that may contribute to variability of choice indices We identified the following factors that may contribute to variability in choice indices in the aversive learning assay across the pilot studies conducted across laboratories (UU, LU, UEA):

1. Light, temperature, and disturbance: Light exposure and environmental temperature affect worm physiology and behaviour and may thereby influence choice indices^50,51^. In our pilot studies, we observed that light exposure and elevated environmental temperature (25-26°C) could stress the worms. Mechanical disturbances, such as moving, shaking or tilting the choice plates could also be stressful for the worms, or present an environmental cue that they perceive and respond to. Thus, all assays included in this manuscript were conducted in the dark and undisturbed under standard temperature (20°C).
2. Transfer method: The method used to transfer worms from training plates to choice plates may affect their choice indices. In our pilot studies, instead of cleaning and transferring worms by washing and pipetting, we manually picked worms from the training plates onto an unseeded plate, allowed them to crawl for 5 minutes to clean their body surfaces, and then picked them onto the choice plates. This method of transferring involved a higher degree of handling and could stress the worms. Therefore, all assays included in this manuscript were conducted using the washing and pipetting method to clean and transfer worms from training plates to choice plates.
3. Choice plate size: Use of larger, 9-cm plates instead of smaller, 6-cm plates as choice plates could affect the concentration of bacterial odours within the confined air space, as well as the distance between bacterial choice spots and the starting point of the worms. Both of which could potentially affect the strength of olfactory cues and the ability of worms to discriminate between the two bacterial sources. Thus, all assays in this manuscript used 6-cm plates as choice plates.

### Implication of the variability of choice indices

An important implication of the sensitivity of the experimental paradigm to environmental conditions concerns the robustness and ecological relevance of transgenerational responses in nature. *Caenorhabditis* worms inhabit highly variable and fluctuating environments in which factors such as light, temperature, food availability, and physical disturbance, constantly change. If transgenerational responses are sensitive to environmental factors under controlled laboratory conditions, they may be even more easily altered under natural condition. This raises the question of whether transgenerational effects observed under controlled laboratory conditions persist under the greater environmental variability experienced in nature.

## Supplemental files

### Supplemental file S1. CERV alignment

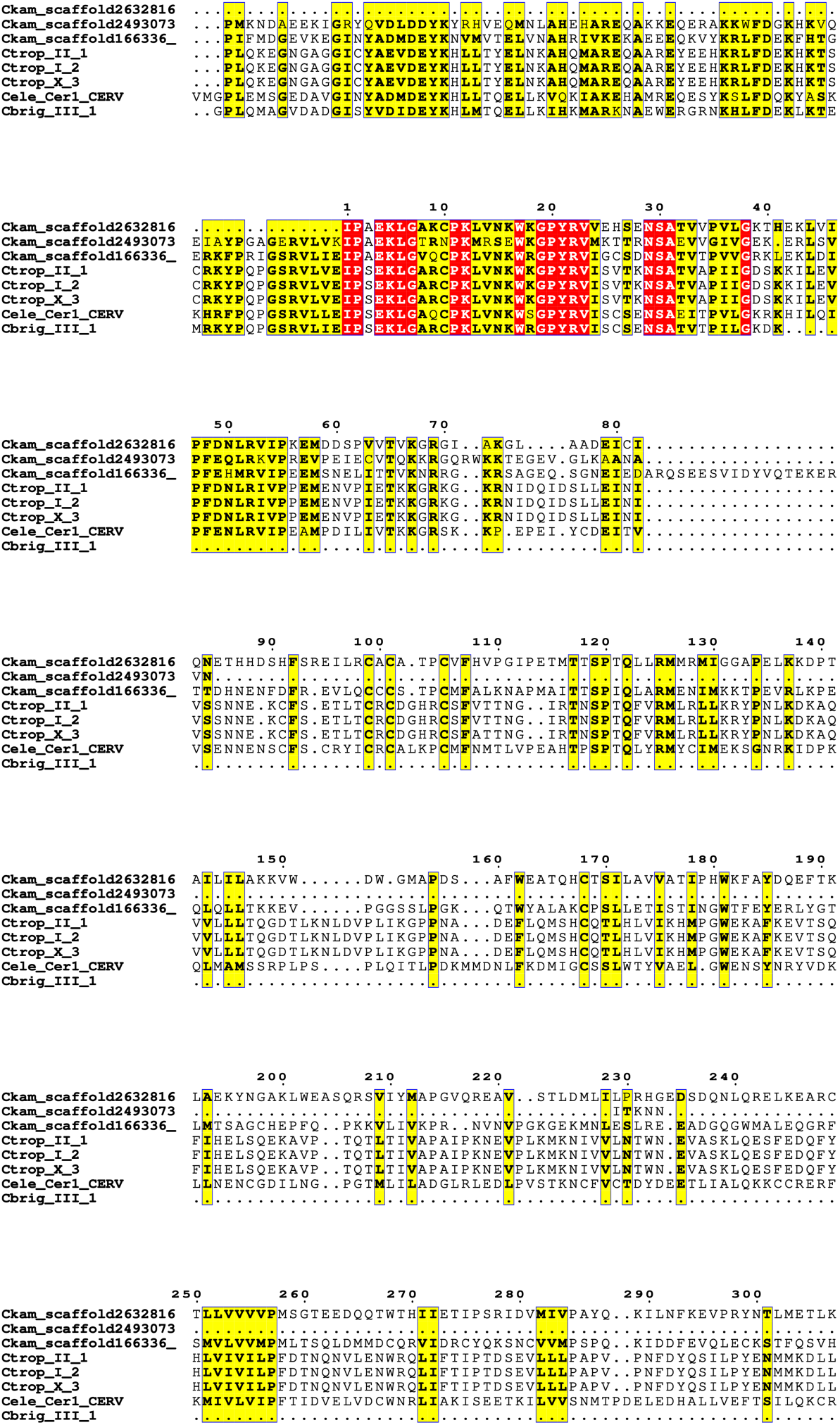

### Supplemental file S2. GAG alignment

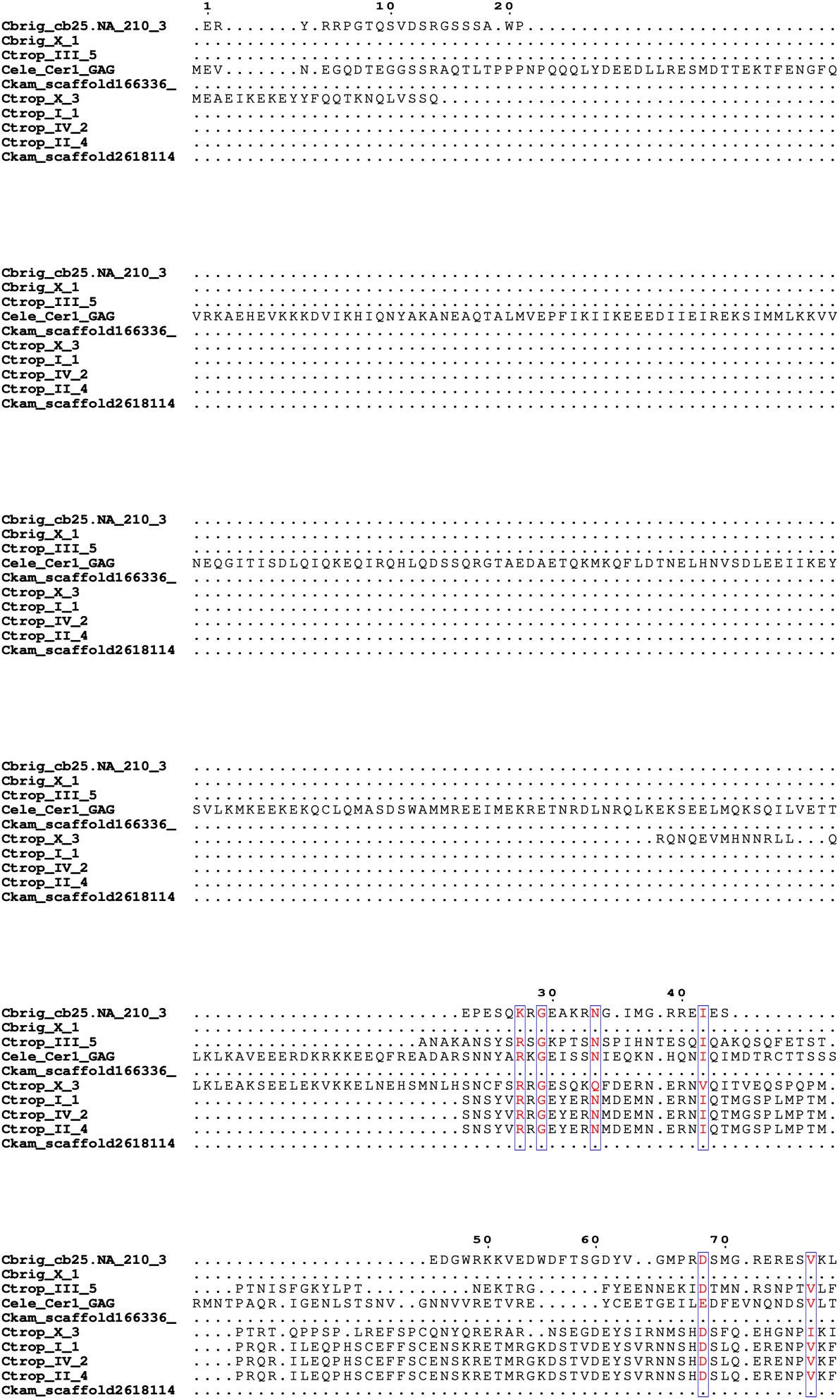

### Supplemental file S3. POL alignment

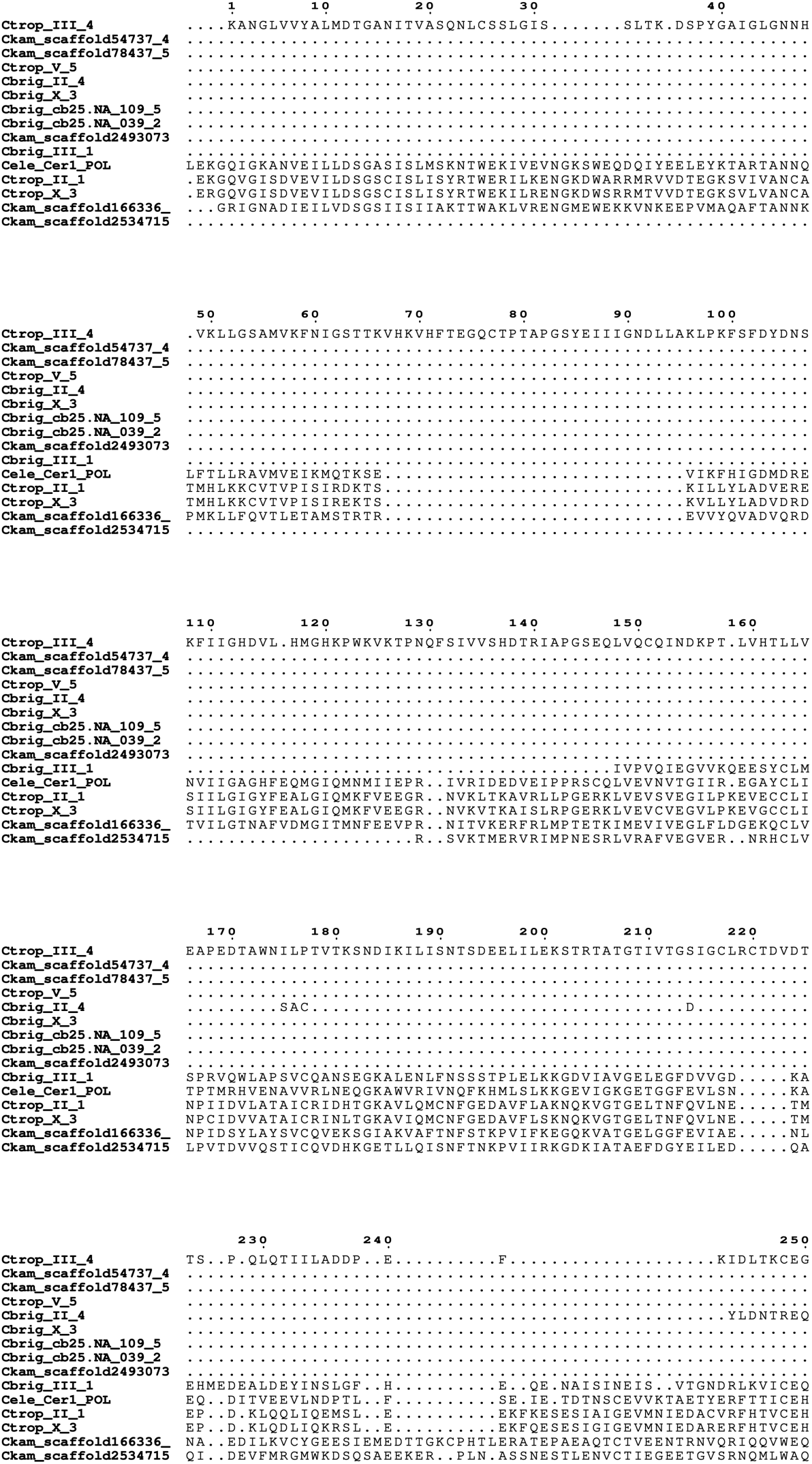

